# Biocontrol potential of grapevine endophytes against grapevine trunk pathogens

**DOI:** 10.1101/2020.09.28.312223

**Authors:** Isidora Silva-Valderrama, Diana Toapanta, Maria de los Angeles Miccono, Mauricio Lolas, Gonzalo A. Díaz, Dario Cantu, Alvaro Castro

**Affiliations:** UC Davis Chile Life Sciences Innovation Center, Providencia, Santiago, RM, Chile; Department of Viticulture and Enology, University of California Davis, Davis, California, USA; Laboratorio de Patología Frutal, Facultad en Ciencias Agrarias, Universidad de Talca, Talca, Chile

**Keywords:** Fungal endophytes and epiphytes, Biological control, Fungal antagonism, Co-culture experiments, GTD

## Abstract

Grapevine Trunk Diseases (GTDs) are a major challenge to the grape industry worldwide. GTDs are responsible for considerable loss of quality, production, and vineyard longevity. Seventy five percent of Chilean vineyards are estimated to be affected by GTDs. GTDs are complex diseases caused by several species of fungi, including *Neofusicoccum parvum, Diplodia seriata*, and *Phaeomoniella chlamydospora*. In this study, we report the isolation of 169 endophytic and 209 epiphytic fungi from grapevines grown under organic and conventional farming in Chile. Multiple isolates of *Clonostachys rosea, Trichoderma* sp., *Purpureocillium lilacium, Epiccocum nigrum, Cladosporium* sp., and *Chaetomium* sp. were evaluated for their potential of biocontrol activity against fungal trunk pathogens. Tests were carried out using two dual-culture-plate methods with multiple media types, including agar containing grapevine wood extract to simulate in planta nutrient conditions. Significant pathogen growth inhibition was observed by all isolates tested. *C. rosea* showed 98.2% inhibition of all pathogens in presence of grapevine wood extract. We observed 100% pathogen growth inhibition when autoclaved lignified grapevine shoots were pre-inoculated with either *C. rosea* strains or *Trichoderma* sp.. Overall these results show that *C. rosea* strains isolated from grapevines are promising biocontrol agents against GTDs.

## Introduction

Grapevine trunk diseases (GTDs) are a major challenge to viticulture worldwide, because they compromise the productivity and longevity of grapevines (*Vitis vinifera* L.) and increase production costs (Munkvold et al. 1994; Bertsch et al. 2013; Kaplan et al. 2016; Gramaje et al. 2018). GTDs are one of the main phytosanitary problems of the grape industry also in Chile (Auger et al. 2004; Díaz et al. 2011). Chile is the first and fourth largest grape and wine exporter in the world, respectively (Felzensztein, 2014; Pizarro, 2018; USDA Foreign Agricultural Center 2019). In 2013, about 22% of the commercial vineyards in Chile showed symptoms of GTDs (Díaz et al. 2013; Guzmán, 2018).

GTDs are caused by fungi that often infect established grapevines through wounds produced during winter pruning (Rolshausen et al. 2010). GTDs can also spread during plant propagation (Aroca et al. 2010; Gramaje and Armengol, 2011), with infections found in dormant wood cuttings and young grafted plants (Gramaje and Armengol, 2011; Waite and Morton, 2007; Billones-Baaijens et al. 2013). In Chile, as in other viticulture areas, the most common microorganisms isolated from arms and trunks of grapevines with symptoms of GTDs are ascomycetous fungi and include *Phaeomoniella* (Pa.) *chlamydospora, Diplodia seriata* De Not., and *Neofusicoccum parvum* (Auger et al. 2004; Díaz et al. 2011; Díaz et al. 2011; Díaz and Latorre, 2013; Besoain et al. 2013).

Currently there are no curative treatments against GTDs beside surgical removal of the infected organs (Surico et al. 2006; Wagschal et al. 2008; Gramaje et al. 2018; Mondello et al. 2018; Sosnowski and Mundy, 2018). GTDs are managed mostly by practices that aim to prevent infections (Gramaje et al. 2018; Mondello et al. 2018). Widely adopted preventive practices include late pruning (Petzoldt, 1981; Munkvold et al 1994), double-pruning (Weberet al. 2007), and the application of protectants on fresh pruning wounds (Díaz and Latorre, 2013). Pruning wounds can be protected by benomyl and tebuconazole (Bester et al. 2007), inorganic compounds as boric acid (Rolshausen and Gubler, 2005), or natural antifungal compounds as organic extracts (Mondello et al. 2018). Manual applications of these formulations as paints are effective, but costly and time-consuming, while spray applications are difficult due to the small surface and orientation of pruning wounds (Bertsch et al. 2013; Rolshausen et al. 2010; Wightwick et al. 2010). In addition, no genetic resistance against GTDs has been found in the grapevine germplasm (Suricoet al. 2006; Wagschal et al. 2008).

Biocontrol of GTDs using microorganisms is a promising alternative. For example, *Trichoderma* spp. are effective as a protectant of pruning wounds (Halleen et al. 2010; Mondello et al. 2018). The goal of our work was to identify microorganisms with biocontrol potential among the natural microbial inhabitants of grapevines. Endophytes are microorganisms that inhabit and colonize the internal plant tissue without causing visible damage or illness in the host (Petrini, 1991; Schulz and Boyle, 2005; Zabalgogeazcoa, 2008). These microorganisms are known to mediate plant-environment as well as plant-pathogen interactions (Zabalgogeazcoa, 2008). The contribution of different epiphytes and endophyte species to plant defenses has been widely documented (Arnold et al. 2003; Azevedo et al. 2000; Pieterse et al. 2014). Plant defense induction and antibiotic substance production that inhibits the growth of pathogens and pests (Mousa and Raizada, 2013), such as fungi (Zabalgogeazcoa, 2008), bacteria (Hardoim et al. 2008), viruses (Lehtonen et al. 2006), and insects (Azevedo et al. 2000) have been reported. The rationale behind focusing on endophytes in the search of effective biocontrol agents against GTDs was two-fold (Wicaksono et al. 2017). First, endophytes are adapted to survive inside grapevines, therefore once applied they should have better chances to establish permanent populations than biocontrol agents selected from other biological systems and therefore provide long-lasting protection (Hardoim et al. 2008; Hardoim et al. 2015; López-Fernández et al. 2016; Zabalgogeazcoa, 2008). Second, endophytes share the same niche with plant pathogens, thus in addition to plant-defense induction and antibiosis, they could also compete for space and nutrients with GTD pathogens (Zabalgogeazcoa, 2008).

Here we report the isolation and identification of endophytic and epiphytic fungi from grapevines grown in commercial vineyards in Chile. From this collection, we selected antagonist candidates and evaluated them for growth inhibition activity against the main GTD fungal species found in Chile, in co-culture, and in planta assays. We provide compelling evidence that endophytic and epiphytic strains of *C. rosea* are strong antagonists of the main GTD species, which makes this species a promising candidate as a biocontrol agent to control GTDs. results.

## Materials and Methods

### 1 Vineyard sampled and plant material

Samples of grapevine (*Vitis vinifera* L.) cv. Cabernet Sauvignon and Chardonnay were collected from four commercial vineyards located in the central valleys in Chile under either organic or, conventional farming systems in May 2017 (**Table 1**). Samples of cv. País were collected in September 2017 from a vineyard where diseases are not managed located in the Codpa Valley, Chile (**Table 1**).

**TABLE 1:** Sample locations

| Vineyard | Variety | Location | Disease control | Planting Year |
| --- | --- | --- | --- | --- |
| Site 1 | Chardonnay | -35°26'26.8764"S, -071°50'01.8600"W | conventional |  |
|  | Cabernet Sauvignon | -35°26'26.8764"S, -071°50'01.8600"W | conventional |  |
| Site 2 | Chardonnay | -34°42'53.3736"S, -071°02'20.5008"W | organic | 2011 |
|  | Cabernet Sauvignon | -34°42'53.3736"S, -071°02'20.5008"W | organic | 2009 |
| Site 3 | Cabernet Sauvignon | -33°44'59.2476"S, -070°56'18.6972"W | conventional |  |
| Site 4 |  | -33°44'59.2476"S, -070°56'18.6972"W | organic | 2000 |
| Site 5 | País | -18°28'42.6"S, -070°05'16.2"W | none | 1850 |

### 2 Isolation of endophytic fungi

The isolation of endophytic fungi was performed following the methodology described in (Pancher et al. 2012). Briefly, shoots (50 cm long) and roots were cut into 10-cm-long fragments. Fragments were surface disinfected by rounds of 2 min serial immersions in 90% ethanol, then 2% sodium hypochlorite solution, and, 70% ethanol, followed by double-rinsing in sterile distilled water under laminar airflow. Absence of microbial growth on surface-sterilized shoots was confirmed by plating the distilled water from the last wash step on potato dextrose agar (PDA; BD-Difco) in Petri dishes, that were then incubated for 2 weeks at 25°C. After disinfection, fragments were further cut into 2.5 mm pieces. Each section was placed on Petri dishes (90-mm diameter), placing the vascular bundle towards the growing media, containing: i) PDA (39 g L^-1^; BD-Difco), ii) malt extract agar (MEA, 33.6 g L^-1^; BD-Difco), and iii) plain agar (AA, 20 g L^-1^; Difco), each one with antibiotics (streptomycin, 0.05 g L^-1^, and chloramphenicol, 0.05 g L^-1^). All Petri dishes were incubated at 25°C for 7 to 10 days under 12 h of light and 12 of darkness. Different colonies were tentatively identified based in morphology (Barnett and Hunter, 1955). Pure cultures were obtained from hyphal tip transfer to PDA media and maintained at 5°C.

### 3 Isolation of epiphytic fungi

For each plant, 1.5 g of soil in direct contact with roots was carefully collected. In a laminar flow bench, 13.5 ml of sterile distilled water was added, before vigorous agitation for 20 min in a horizontal position. After 5 min of decantation, serial dilutions of the supernatant were made. 10^-3^ and 10^-4^ dilutions were used to inoculate PDA, MEA, and AA. To all media streptomycin, 0.05 g L^-1^ and chloramphenicol, 0.05 g L^-1^ were added. Plates were incubated for 7 to 14 days at 25°C.

### 4 Taxonomic characterization of the fungal isolates

DNA extraction from cultivable isolated fungi (n=387 isolates) was performed as described in Morales-Cruz et al. (2015), with the following modifications. Mycelia from 7 to 21 days old fungal cultures were frozen with 3 mm metal beads in tubes at −80°C. Tubes were shaken vigorously with a vortex for 5 minutes at maximum speed. Disrupted mycelia were resuspended in 200 μL of nuclease-free sterile-distilled water and then homogenized in a vortex for 15 s. Mycelia was incubated at 100°C for 10 min, followed by a centrifugation step at 14500 rpm for 2 min. An aliquot of 10 μL of the supernatant was used for the PCR runs. A 1:20 or 1:50 dilution was made in case of PCR inhibition occurred. ITS sequences were PCR amplified using ITS1 (TCCGTAGGTGAACCTGCGG) and ITS4 (TCCTCCGCTTATTGATATGC) primers (White et al. 1990). A 25 μL PCR reaction was carried out using 2.5 uL 1XThermopol reaction buffer, 0.5 uL of 10mM dNTPs, 0.5 uL of 10uM ITS forward and reverse primers, 0.125 μL (1.25U/50 μL) Taq DNA polymerase (Promega, USA) and 10 μL of sample supernatant as a template. PCR reaction was performed with an initial denaturing step at 95 °C for 2 min, and 35 cycles of 95 °C for 30 s, 52°C for 30 s (White et al. 1990), and 72°C for 1 min, followed by a final extension phase at 72 °C for 5 min. The PCR product was purified and sequenced at Macrogen Inc., South Korea. Amplicon sequencing analysis was carried out with Geneious (R11.1). Taxonomic identities were determined with BLASTN using the UNITE database 7.2 (Nilsson et al. 2019).

### 5 Pathogenic fungal strains and control antagonists

Isolates of *Phaeomoniella chlamydospora* (#11 A), *Diplodia seriata* (N°117 Molina), *Neofussicoccum parvum* (N°156 Lolol) and the endophytic antagonist *Trichoderma* sp. (Altair 607 QR6 PB 6.0) were obtained from the Phytopathology Lab of Universidad de Talca. These isolates were purified in 2017 from *V. vinifera* L. trunks as part of another project. Also, MAMULL (*Trichoderma gamsii* Volqui strain, *Bionectria ochroleuca* Mitique strain, *Hypocrea virens* Ñire strain, BioInsumos Nativa, Chile), TIFI (Giteniberica de Abonos, España), Tebuconazole 430 SC (SOLCHEM, concentrated suspension, Chile) were used as positive controls.

### 6. Test of fungal antagonism

Initial assessment of antagonistic properties was conducted against *D. seriata* as pathogen. Further evaluations on selected antagonists were carried out using *D. seriata, N. parvum*, and *P. chlamydospora*. Agar discs from a 7-day old actively growing colony were used. Co-culture assays were performed placing a 5 mm agar disc on one side of the Petri dish with PDA (39 g L^-1^; Difco) or PA (200 g L^-1^ grapevine propagation material, 20 g L^-1^ agar) and on the opposite side a 5 mm agar disc containing the antagonist strain. Plates were incubated at 25°C for 7-28 days in darkness (Badalyan et al. 2002) using a randomized complete block design. Registered bioproducts MAMULL and TIFI were used as antagonistic controls. Pathogen growth area was evaluated at 7, 14, 21, and, 28 days post-co-culture (Schindelin et al. 2012). Inhibition percentage was calculated using the pathogen growth area when was cultured alone (C) or in interaction with the antagonist (T) according to the formula I = ((C-T)/C) * 100 (Thampi and Bhai, 2017).

An in planta assay was also performed. Annual shoots were used for the experimental set-up to verify the antagonistic potential shown in plate co-culture. Several preliminary evaluations were carried out in order to test variability caused by autoclave sterilization of pruning material, humid-chamber moist maintenance, type of inoculum and time needed for the pathogen to grow through the wood piece. Even though tissue was death, the overall shoot matrix structure was conserved after autoclave sterilization (data not shown). Internode portions of dormant cuttings were cut in 4.5 cm length pieces and then used fresh or autoclaved for 25 min at 121 °C. Agar mycelium plugs were evaluated as inoculum. In 2 days, pruning material in contact with the pathogen and/or antagonist plugs were covered in the mycelium. As the inoculum was too high, a spore suspension solution was used to inoculate the wood pieces. Mycelium/spore mix suspension of the pathogens *D. seriata* and *N. parvum* were prepared by flooding 30 days old plant agar culture (PA; 200 g L^-1^ grapevine dormant cutting, 20 g L^-1^ agar) with sterile distilled water. In the case of the antagonists *Clonostachys rosea* (isolates CoS3/4.24, CoR2.15 and R31.6) a spore suspension adjusted to 1 x 107 conidia mL^-1^ was used as recommended. Antagonist inoculation was carried out adding 40 uL of antagonist fresh spore suspension until it reached the woody stem cut end by capillarity. Tebuconazole (60 mL/100L fields recommended doses; SOLCHEM, Chile) or sterile distilled water was applied in the same manner as controls. This experiment was carried out 5 times. Woody stem cuts were incubated in individual humid chambers for 24 hours. Then, 10 uL of fresh pathogen mycelia/spore mix suspension was inoculated on the same side where the antagonist was inoculated previously and immediately placed in a horizontal position, preventing suspension diffusion. Incubation was carried out in humid chambers for 3-7 days. Afterward, the surface of the woody stem was disinfected by rubbing with 70% ethanol. With a hot sterile scalp, the bark and 0.5 cm of the woody stem ends were removed. Small pieces located at 1 and 2.5 cm from the inoculation point were collected and cultured in individual PDA plates at 25°C for 7 days. To evaluate the pathogen mycelia and spore suspension viability, 10 uL of the solution was inoculated in one side of the wooden piece as described above and immediately processed to obtain 3 mm pieces at 1 and 2.5 cm from the pathogen inoculation point. Every piece was cultured in PDA at 25 °C for 7 days. The presence of the pathogen on PDA was evaluated under a light microscope.

### 7 Test of antagonist mechanism

To characterize the mechanism of antagonism, the same experimental setup of co-culture was carried out on water agar (AA, 20 g L^-1^; Difco) with a microscope sterile slide covered by a thin layer of the same agar in its surface. Using a light microscope (MOTIC BA410), the sample was screened for loops of the antagonist hyphae around *N. parvum* and *D. seriata*, indicating mycoparasitism. This experiment was carried out 3 times. To determine antibiosis as the type of antagonist mechanism used, isolated fungi *E. nigrum* R39.1, *C. rosea* CoS3/4.4, and *Cladosporium* sp. B38d.2 were cultured in PDA plates (39 g L^-1^; Difco) over cellophane paper for 7 days. Cellophane paper with the fungal colony was then removed from the plate and a mycelial plug of *D. seriata* or *N. parvum* was placed in the centre. Plates were incubated for 7 days at 25°C and pathogen growth was evaluated. This experiment was carried out three times.

### 8 Statistical analysis

Statistical analysis was conducted with GraphPad PRISM 8 (8.1.1 version, 2019).

## Results

### 1. Isolation and identification of endophytic and epiphytic fungi

A total of 102 vineyard samples were collected to isolate endophytic and epiphytic fungi associated with grapevines in Chile. Endophytic fungi were isolated from woody shoots, sprouts, and roots, while the epiphytic ones were obtained from the rhizosphere. Ninety samples were obtained from two commercial vineyards in the central valleys of Chile and twelve from a vineyard in the Codpa Valley that has not been managed for disease protection for over 150 years. A total of 222 and 166 morphologically distinct filamentous fungi and yeasts were isolated from the commercial vineyards and the Codpa Valley, respectively. Fungi were isolated and characterized taxonomically using ITS1 and ITS4 sequences. All fungal sequences were at least 98% identical to the best BLASTn hit in the UNITE database. We could assign taxonomy to a total of 300 isolates. The ITS sequence was discriminant at the species level for 227 isolates. The remaining were assigned to the corresponding genus or family. A total of 58 genera were represented, 37 and 38 among epiphytes and endophytes, respectively. As expected, below ground samples (rhizosphere and roots) were more diverse (56 genera) than sprouts and woody stems (5 genera) (**Figure 1**).

**FIGURE 1.**
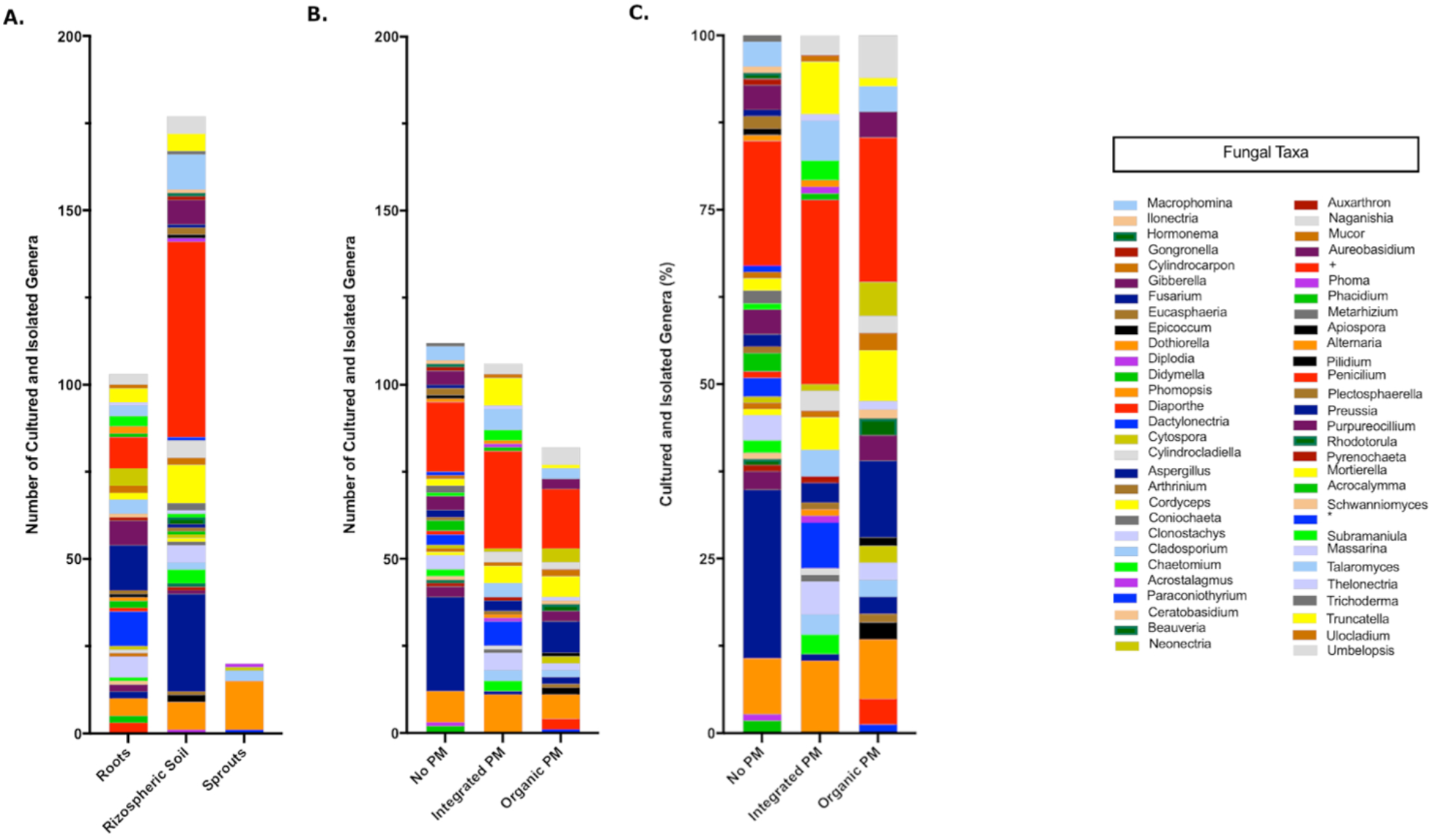
Taxonomic composition of the isolated fungi. Values are separated according to the source (A) and phytosanitary regime (pest management program, PM) (B)(C). Cultured-isolates identified only to family level Nectriaceae (+) and class level Dothideomycetes (*) are also shown.

### 2. Effect of fungal antagonists on the growth of GTD fungi in co-culture

To identify potential biocontrol agents for further characterization, we screened all isolates for antagonistic activity against *D. seriata* (**Supplementary Table S1**), a ubiquitous GTD pathogen. Based on the results this initial screen, a total of ten isolates were selected for further characterization: *Trichoderma* ap. Altair, *Epicoccum nigrum* R29.1, three isolates of *Clonostachys rosea* (R 31.6, CoR2.15 and CoS3/4.24), *Cladosporium* sp. B38d.2, *Chaetomium* sp. S34.6 and *Purpureocillium lilacium* S36.1. These ten isolates were chosen also because they were previously described as antagonistic to other pathogens (Fávaro et al. 2012; Hung et al. 2015; Cota et al. 2009; Solano Castillo et al. 2014; Costadone and Gubler, 2016).

To assess the antagonistic ability of the ten selected isolates, we co-cultured each one of them with *D. seriata* and *N. parvum*, two of the main fungi causing GTDs in Chile. Co-cultures were carried out on two different types of growth media: the commonly used potato dextrose agar (PDA) and a substrate made of agar and ground woody grapevine tissue (aka, grapevine plant agar (PA)) that simulates in planta nutrient composition (Massonnet et al. 2017). Isolates displayed a wide range of growth rates, which often differed between PDA and PA (**Figure 2**). Interestingly, most endophytes, including all *C. rosea* isolates, grew faster on PA than PDA. Different growth rates reflected the patterns of inhibition of *D. seriata* and *N. parvum* (**Figures 3 and 4**). The *Trichoderma* Altair isolate grew faster than the rest on PDA and reached its maximum inhibitory effect on both pathogens as early as day 7 in PDA. Growth inhibition only occurred upon physical contact between colonies of *Trichoderma* sp. and the pathogens. The faster growth on PA of the endophytes *Clonostachys, Chaetomium, Epicoccum*, and *Cladosporium* was associated with greater pathogen inhibition rates on this substrate compared to PDA, especially for the *Clonostachys* isolates. In PA, *C. rosea* overgrew the pathogen colony at least 7 days earlier than in PDA. All *C. rosea* strains inhibited over 98% pathogen growth in PA at day 21 (**Figure 4**). *Chaetomium* sp. S34.6 isolate inhibited pathogen growth by slowly growing in the plate until colony contact. By day 21 *Chaetomium* sp. S34.6 inhibited *D. seriata* and *N. parvum* growth by 59.1% and 86.75%, respectively, about two-fold the pathogen growth inhibition showed in PDA. Both species completely overgrew both pathogen colonies around 28 days. The antagonistic effect of *C. rosea* R36.1 and CoS3/4.24 occurred upon direct contact between colonies, which overgrew the pathogen colony within 21 days of growth. Instead, pathogen growth inhibition of *C. rosea* CoR2.15, *Purpureocillium lilacium* S36.1, and *E. nigrum* R29.1 happened without evident physical contact between colonies. In PDA, *E. nigrum* produced a wide 0.8 to 1.2 cm orange-colored halo that was partially colonized only by *N. parvum* after 21 days of growth. The slow and limited growth of *Neofusicoccum parvum* was also visible in the halo produced by *Purpureocillium. Cladosporium* sp. B38d.2 showed an interesting difference in antagonist activity against N. parvum in PA, reaching its higher inhibition rate (**Figure 4**). When cultured with this pathogen*, Cladosporium* strongly sporulated, covering the entire plate, and stopped *N. parvum* early growth.

**FIGURE 2.**
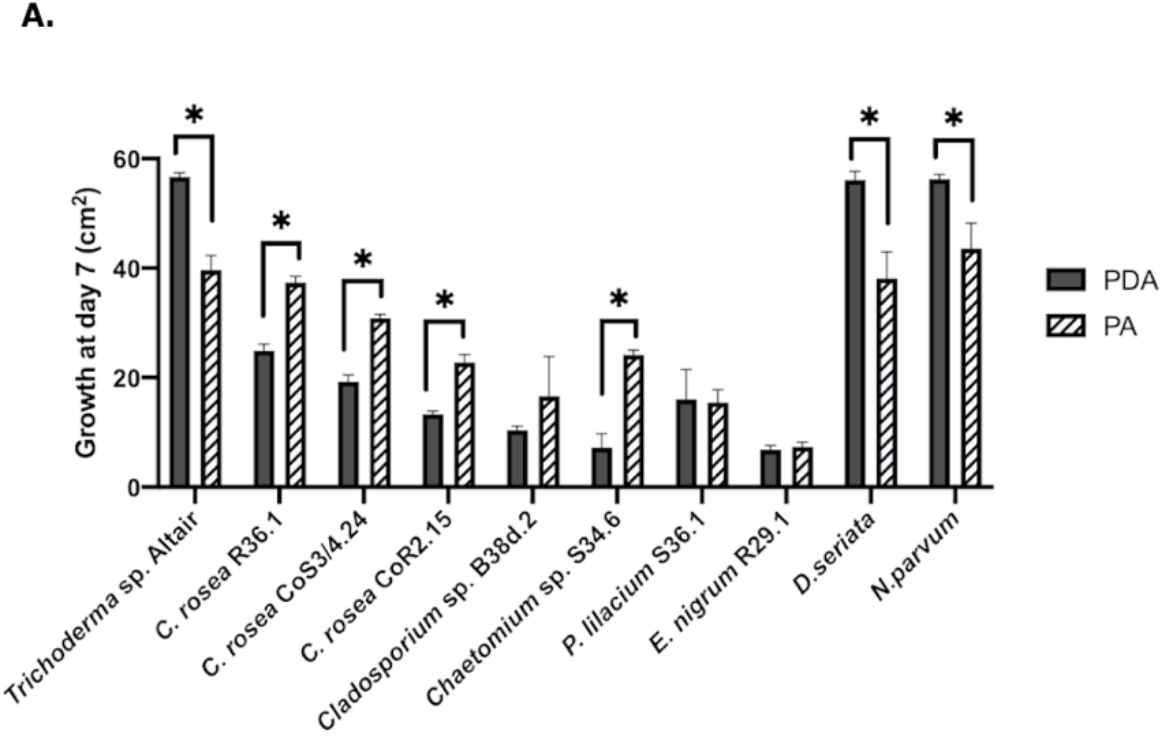
Comparison of the growth area of antagonists and pathogens in two media. Growth was measured after 7 days in PDA (potato dextrose agar) and PA (plant agar). Bars with asterisk are significantly different from the control (Paired T test, P<0.001). Error bars represent the standard error of the mean, n=5.

**FIGURE 3.**
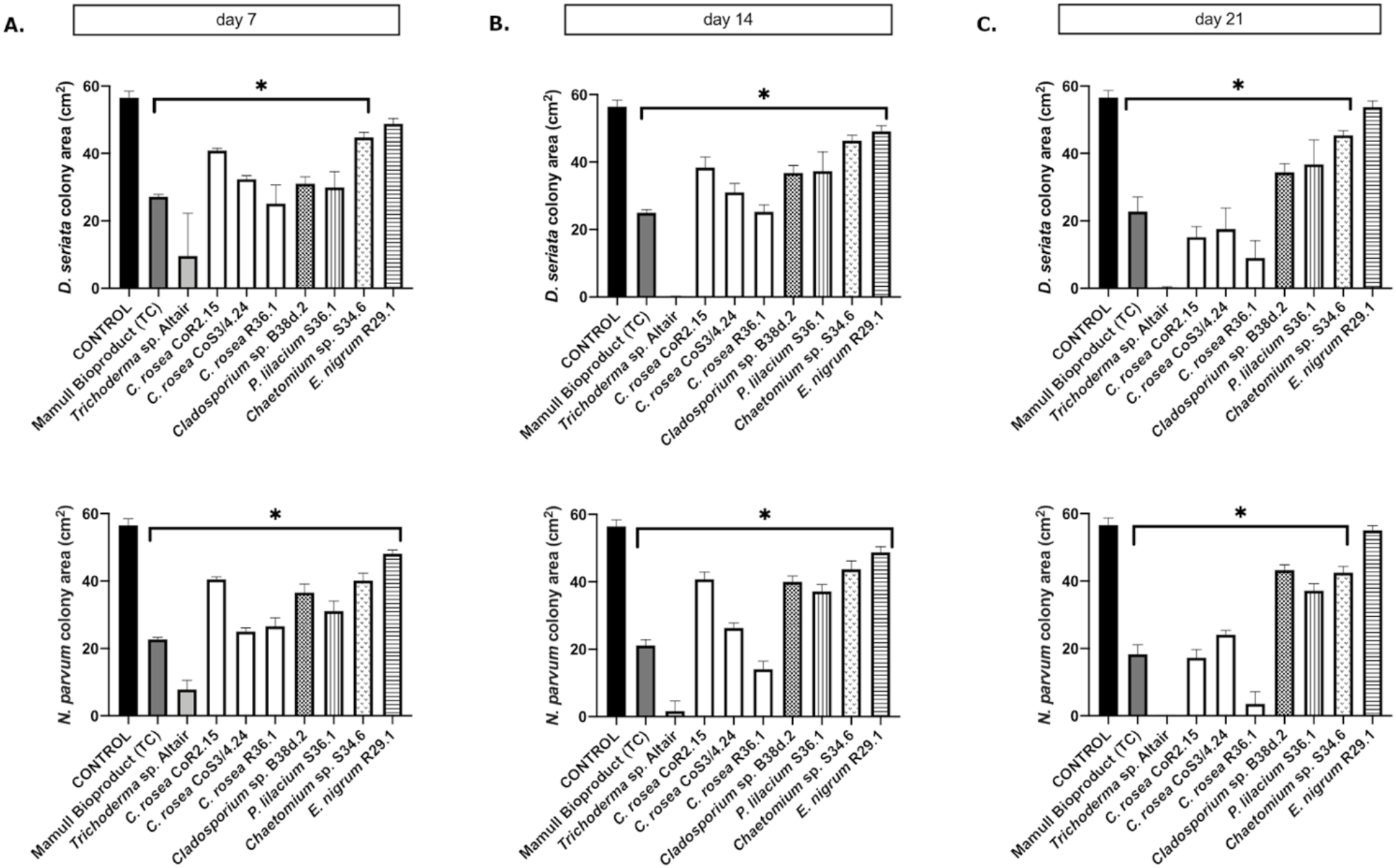
Colony area measured after (A) 7, (B) 14 and (C) 21 days of inoculation of *D. seriata* (upper graphics) and *N. parvum* (bottom graphics), when growing alone (control) or in co-culture with the antagonists in PDA. Bars with asterisk are significantly different to the control (Tukey’s test, P<0.001). Error bars represent the standard error of the mean, n=5.

**FIGURE 4.**
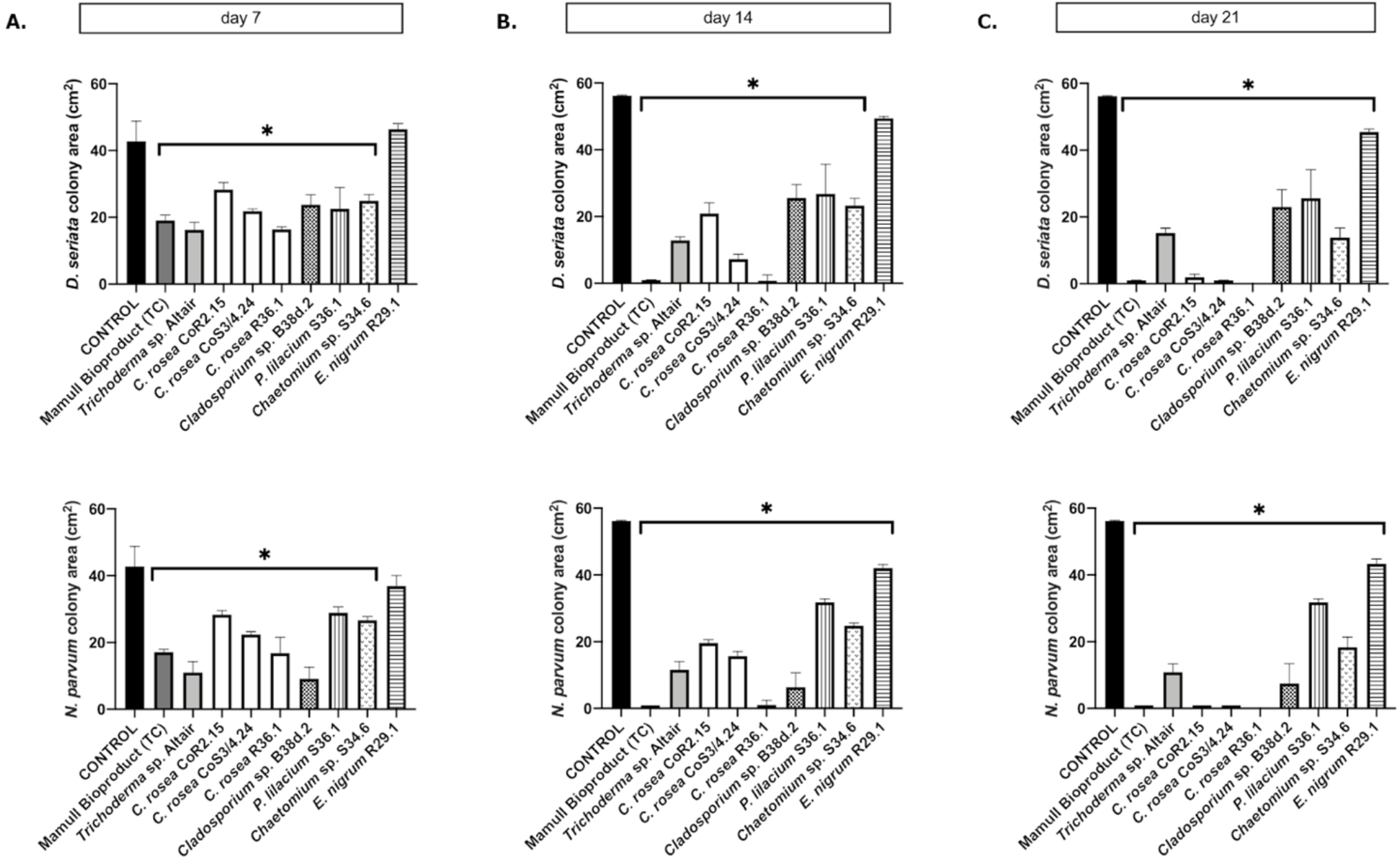
Colony area measured after (A) 7, (B) 14 and (C) 21 days of inoculation of *D. seriata* (top row) and *N. parvum* (bottom row), when growing alone (control) or in co-culture with the antagonists in PA. Bars with asterisk are significantly different to the control (Tukey’s test, P<0.001). Error bars represent the standard error of the mean, n=5.

On PA, *C. rosea* inhibited *P. chlamydospora* almost completely (99.9%). Interestingly, *C. rosea* growth first paused without evident contact between colonies (**Figure 5**) at day 7, but later, by 14 days, it overgrew completely the pathogen colony. Overgrowth was also observed with *Trichoderma* sp. Altair in PDA.

**FIGURE 5.**
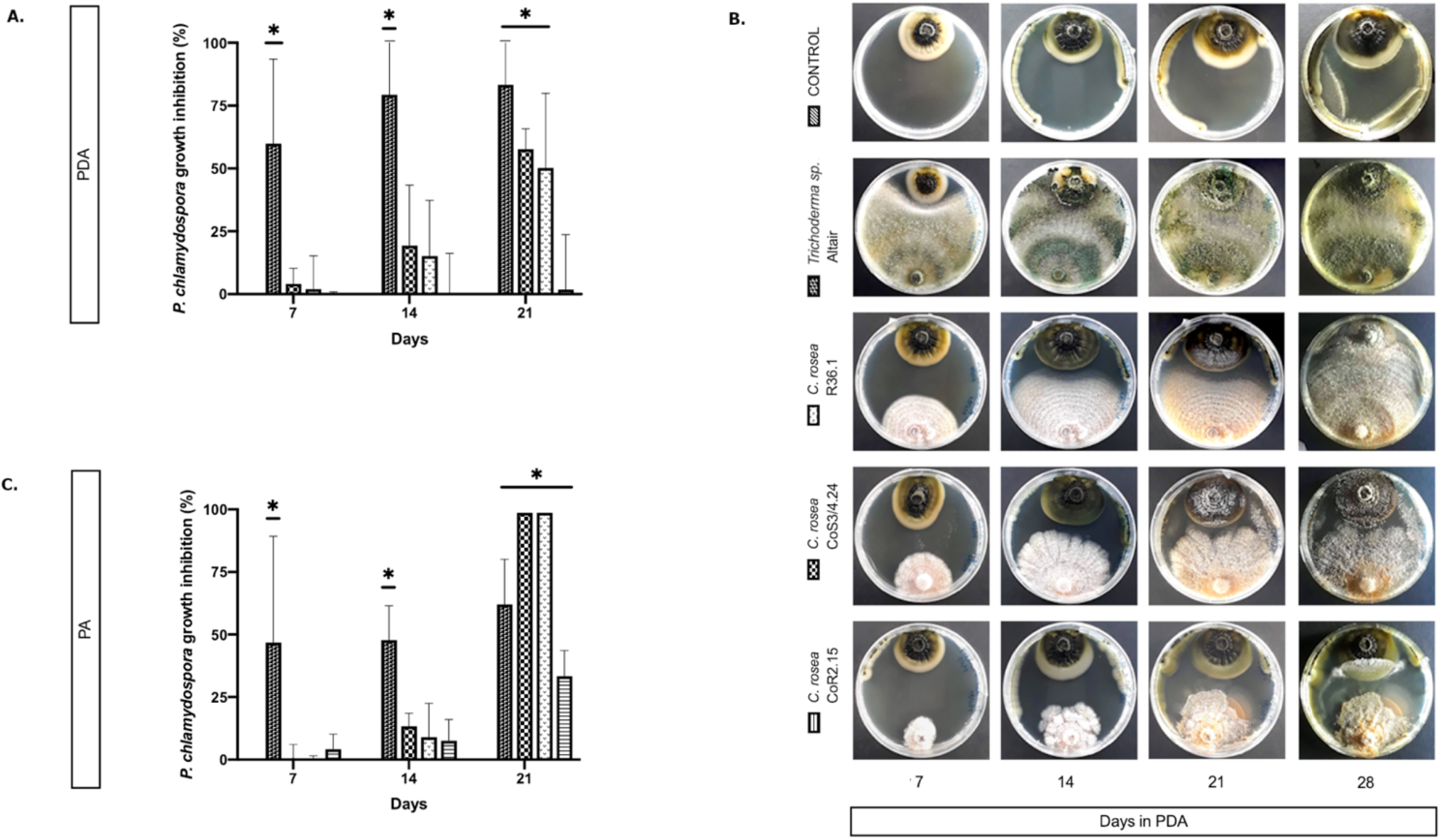
Colony area measured at day 7, 14 and 21 postinoculation of the pathogen *P. chlamydospora* when cultured alone or with the antagonists: *C. rosea* CoR2.15, CoS3/4.24, R36.1 or *Trichoderma* sp. Altair. Growth area was evaluated in potato dextrose agar, PDA, (A) and (B) and, in grapevine plant agar, PA, (C). Bars with asterisk are significantly different to the control (Tukey’s test, P<0.001). Error bars represent the standard error of the mean, n=5.

### 3. Characterization of the mechanisms of antagonism

The antagonistic activity of endophytic biocontrol agents can depend on the competition for nutrients and induced resistance in the plant, and/or direct interaction with the release of pathogen inhibitory compounds or mycoparasitism (Köhl et al. 2019). During co-culture, isolates of *C. rosea* showed pathogen inhibition both before and after direct contact between colonies, suggesting that both mechanisms could underlie its antagonistic properties. To evaluate the mode of action of *C. rosea* and *Trichoderma* sp. Altair, we studied under a light microscope the mycelia in the zone of interspecific interaction. For *C. rosea* CoS3/4.24 and R36.1, hyphal coiling, a sign of mycoparasitism, was consistently observed in all co-cultures with *N. parvum* and *D. seriata* (**Figure 6**). Hyphal coiling was only occasionally found in *Trichoderma* sp. Altair.

**FIGURE 6.**
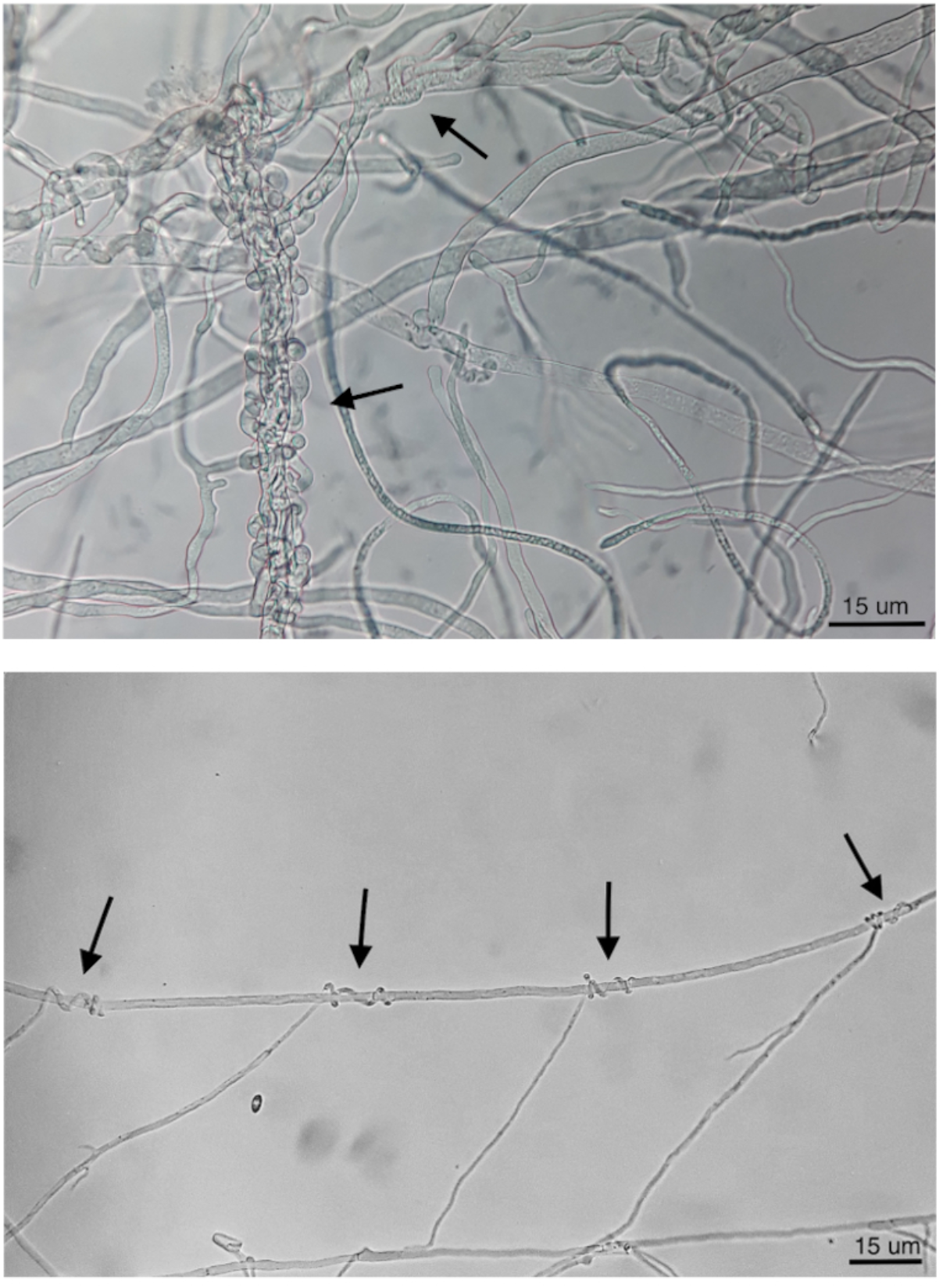
Hyphal coiling of (A) *Trichoderma* Altair against *D. seriata* and (B) *C. rosea* CoS3/4.24 around hyphae of *N. parvum* (magnification 400X).

When *C. rosea* epiphytic strain CoS3/4.24 was co-cultured with *D. seriata* or *N. parvum*, pathogen growth terminated before direct contact with *C. rosea* in correspondence of the halo surrounding the antagonist. In this case, the inhibitory activity of *C. rosea* may depend on a secreted antibiotic compound. This was also observed when *Cladosporium* sp. B38d.2 was used as antagonist. To test the inhibitory activity of the *C. rosea* secretome, we inoculated *C. rosea* on a sterilized cellophane membrane overlaid on PDA and incubated for seven days. The cellophane membrane was shown to be permeable to metabolites secreted by fungi (Dennis and Webster, 1971; Chambers, 1993; Sharmini et al. 2004; Rodriguez et al. 2011). After removing the cellophane membrane together with the *C. rosea* mycelia, we inoculated the plates with pathogens and measured their growth in comparison with normal PDA. Pathogen growth was significantly reduced on plates previously incubated with *C. rosea*, likely due to the secreted metabolites that permeated through the cellophane membrane (**Figure 7**). The inhibition caused by the secreted metabolites of *C. rosea* CoS3/4.24 led to a 47.2% and 50.1% reduction in growth of *D. seriata* and *N. parvum*, respectively. In the case of *Cladosporium* sp., 34.26% and 42.46% inhibition was observed against *N. parvum* and *D. seriata*, respectively. Changes in the pathogen colony morphology were also observed, especially when in contact with *C. rosea* CoS3/4.24 isolate secondary metabolites. *N. parvum* colony turned into several flat independent colonies with undulate margins, while *D. seriata* grew as one colony with irregular shape.

**FIGURE 7.**
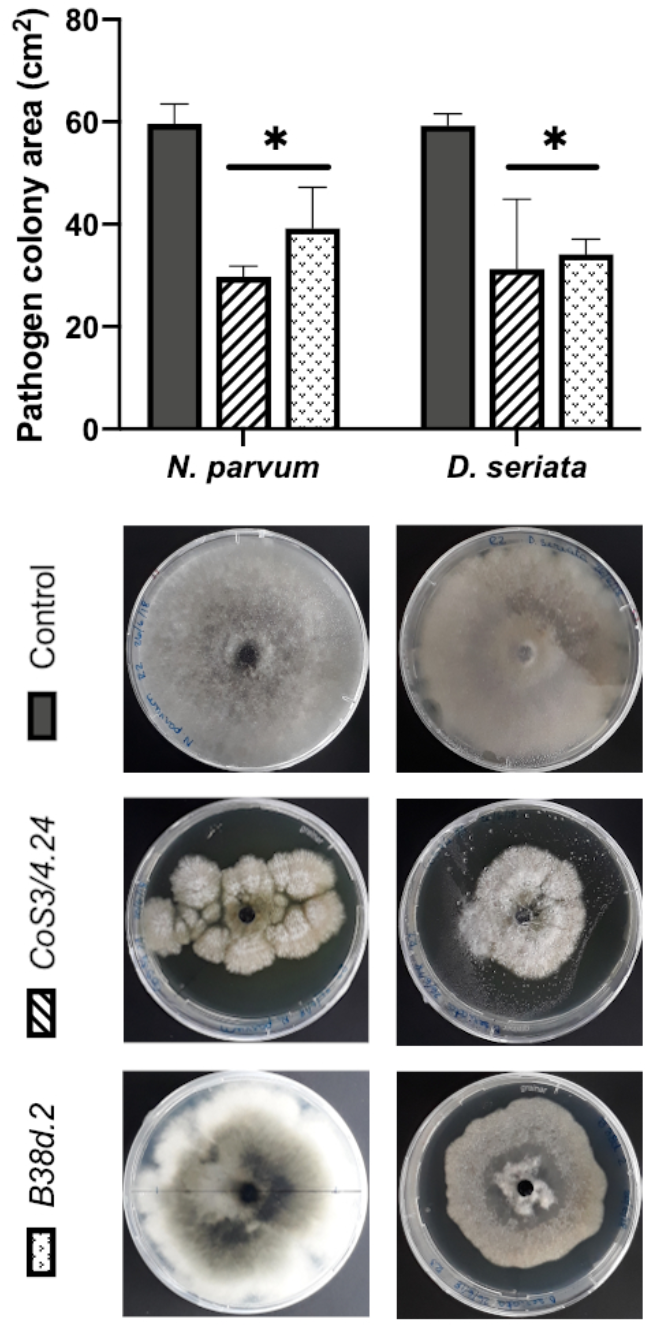
Pathogen growth over secondary metabolites produced by antagonists *C. rosea* CoS3/4.24, *Cladosporium* sp. B38d.2 in PDA. Bars with asterisk are significantly different to the control (Tukey’s test, P<0.001). Error bars represent the standard error of the mean, n=5.

### 4. Effect of fungal antagonists on the growth of GTD fungi in one-year old grapevine woody shoots

As both growth and inhibition rates of GTD pathogens were significantly different in media containing grapevine annual shoot extract (plant agar, PA), we extended the testing of antagonism by using one-year-old lignified shoots (aka canes) as a substrate for co-cultures. We tested both sterile (autoclaved) and non-sterile canes. After 7 days, *C. rosea, N. parvum*, and *D. seriata* colonized completely the internal tissue of 4.5 cm-long autoclaved canes. The antagonists *C. rosea* strains were recovered in all pathogen co-inoculated samples after 7 days (**Figure 8**). No pathogen growth was observed at 0.5 cm from the pathogen inoculation point when treated with the antagonists. Interestingly, under the same conditions, Tebuconazole, a commercial synthetic fungicide, did not reduce *D. seriata* nor *N. parvum* growth.

**FIGURE 8.**
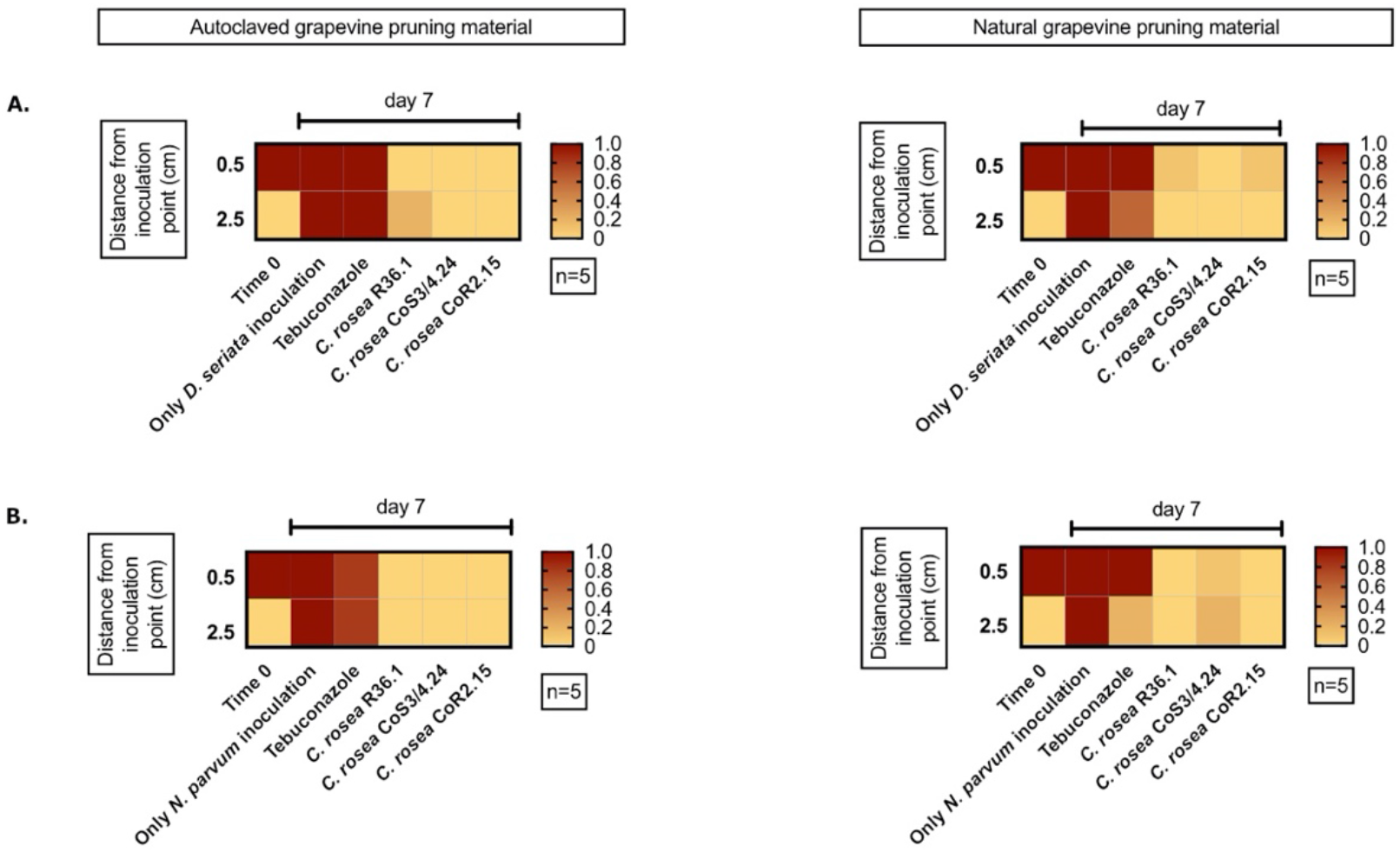
Presence of the pathogen *D. seriata* (A) and *N. parvum* (B) in autoclaved (left graphics) and natural (right graphics) grapevine pruning material pre-inoculated with the antagonist. In red is shown 100% recuperation of the pathogen.

We also performed the co-culture experiments on canes that were not subjected to autoclaving. Pathogens colonized the entire cane in 7 days in absence of any antagonist. In less than 0.1% and 10% of the co-culture assays, *N. parvum* and *D. seriata* were recovered from plant tissue previously inoculated with *C. rosea* isolates, respectively. In the case of CoS3/4.24 isolate, *N. parvum* and *D. seriata* growth inhibition was observed in 80% and 100% of the assays, respectively. In summary, the antagonistic potential of the *C. rosea* isolates shown in agar plate was confirmed in grapevine propagation material.

## Discussion

We isolated fungi from grapevines to find potential biocontrol agents against GTDs. As they share the same host with pathogens, these fungi may provide longer-lasting protection of grapevine tissues than biocontrol agents identified on other plant species (Zabalgogeazcoa, 2008; Latz et al. 2018). Three hundred eighty-seven different fungi and yeast were isolated and identified from multiple grapevine tissues and pest management systems. The observed diversity was limited to culturable fungi, since no cultivation-independent identification tools were applied. Taxa were determined solely based on the ITS sequence. Further validation using other informative sites, such as nu-SU-0817-59 and nu-SU-1196-39 (Borneman and Hartin, 2000) or TEF-1a (Ichi-Ishi and Inoue, 2005), would provide additional resolution for some of the isolates we were not able to characterize at the species level. As expected, rhizospheric soil showed to hold more fungal diversity than roots, and sprouts showed less cultivable diversity than any other sample. This was in agreement with previous studies using amplicon sequencing (Tan et al. 2017).

As the focus of this work was to find microorganisms able to colonize the grapevine persistently, we conducted this search during late Winter, at the beginning of the cold and wet season, when potentially beneficial microorganisms may compete with pathogens for the colonization of the host through pruning wounds (Rolshausen et al. 2010; Travadon et al. 2016; Arnold et al. 2003). Even if we could collect more samples from commercial vineyards than from the 150 year-old vines in the Codpa valley, the number of fungal taxa isolated from Codpa was higher than in commercial vineyards. The greater diversity found in Codpa might be due to the older age of the vines as well as the lack of pathogen control practices throughout the life of the vineyard, even if other cultural management practices as fertilization with animal manure have been done over generations.

All fungi we isolated, characterized, and tested, with the exception of *Epicoccum nigrum* showed a significant growth inhibition of *N. parvum* and *D. seriata* in co-cultures on both PDA and PA. The *Trichoderma* Altair isolate and all *C. rosea* strains completely overgrew both pathogens by day 21. This was also observed against the pathogen *P. chlamydospora* in PA. However, variable biocontrol efficacy was observed between different isolates of the same species, as reported in (Inch and Gilbert, 2007). For example, the epiphytic isolate *C. rosea* CoS3/4.24 grew faster on media and overgrew the pathogen earlier than the other *C. rosea* isolates. In contrast, the endophytic isolates of *C. rosea* showed better inhibition of *N. parvum* in grapevine woody shoots. The endophytic isolate of *Cladosporium* also displayed antagonism in co-culture, in particular against *N. parvum* on PA. Its inhibitory activity seemed to be due to the high sporulation rate and not to the rapid growth of the mycelium observed in others (Schöneberg et al. 2015). *Cladosporium* sp. produces a great amount of black, hydrophobic spores, and a small mycelium underneath the dense spore mass. On PA as well as PDA, *Chaetomium* sp. showed a significant reduction of growth of *N. parvum* and *D. seriata*, although weaker than that of *Trichoderma*. The antagonistic activity of *Chaetomium* may be due to a slow mycoparasitism. Hyphae of *Chaetomium* has been described to penetrate and coil around pathogen hyphae at day 30 of co-culture (Hung et al. 2015). Strains of *Chaetomium* have also shown antagonist activity against different pathogens as Phythophthora nicotianae (Hung et al. 2015), *Rhizoctonia solani* (Gao et al. 2005) and *Fusarium oxysporum* (Huu Phong, 2016) among others. Some strains presented antibiosis as an antagonist strategy, but mycoparasitism has been also described for this genus (Hung et al. 2015).

*C. rosea* showed limited antagonism at early stages of co-culture on artificial media and completely inhibited pathogen growth only after 21 days. Importantly, *C. rosea* was particularly effective against pathogen colonization of autoclaved woody shoots. Fungal growth dynamics and, therefore, the interaction between colonies are likely influenced by the type of media (Schöneberg et al. 2015b), in particular when nutrient-rich media are compared with substrates poor in nutrients, such as PA and woody tissue. It is worth noting that different isolates displayed different antagonistic activities depending on the substrate. For example*, C. rosea* isolates R36.1 and CoR2.15 showed higher pathogen inhibition than CoS3/4.24 on woody shoots that were not autoclaved. Interestingly, R36.1 and CoR2.15 were endophytic, while CoS3/4.24 was isolated from the rhizosphere. Although we did not find the same pattern when autoclaved tissue was used, the different behavior of endophytic and epiphytic isolates supports the overall strategy to search for potential biocontrol agents among the natural inhabitants of grapevines.

Generally recognized control mechanisms for fungal biocontrol agents are (1) competition for nutrients and space, (2) induced resistance in the plant, both consisting in an indirect interaction with the pathogen, (3) inhibition through antibiosis, and, (4) mycoparasitism (Latz et al. 2018; Köhl, 2019). The formation of short loops of the antagonist’s hyphae around hyphae from another fungal species also called hyphal coiling (Assante et al. 2004; Barnett and Lilly, 1962; Gao et al. 2005). The coiling establishes an intimate contact with the parasitized hypha, penetrating the hypha and delivering antibiotic compounds and cell-wall degrading enzymes (Barnett and Lilly, 1962). This type of mycoparasitism has been commonly found in the genus *Trichoderma* (Howell, 2003; Benítez et al. 2004) and reported in *C. rosea* (Barnett and Lilly, 1962; Morandi, 2001). The Trichoderma sp. Altair isolate produced hyphal coils and also the *C. rosea* strains we tested. In all cases, we found a strong correlation between coiling and antagonism suggesting that mycoparasitism plays an important role in the interaction with the pathogens. In the case of *C. rosea* CoS3/4.24, a yellowish halo around the antagonist colony was present. Antibiosis was previously described for this species (Iqbal et al. 2017), but not all strains of the species show antibiotic production (Moraga-Suazo, 2016). Further studies should be performed with the *C. rosea* isolates as this might have important applications in agro-industrial areas (Karlsson et al. 2015). Direct interaction with the pathogen mode of action, as mycoparasitism and antibiosis, are highly desirable mechanisms for further production of commercial biocontrol agents, as they expose lower risks of human, plant and, environmental toxicity (Köhl, 2019).

## Acknowledgments

We would like to thank the VSPT group for let us collect samples from their vineyards and supporting this project. We would also like to thank Patricio Muñoz for providing samples of old grapevines from the Codpa valley.

## Conflict of Interest

No conflict of interest declared.

## Notes

### Competing Interest Statement

The authors have declared no competing interest.

## References

Arnold, A. Elizabeth, Luis Carlos Mejía, Damond Kyllo, Enith I Rojas, Zuleyka Maynard, Nancy Robbins, and Edward Allen Herre. (2003) “Fungal Endophytes Limit Pathogen Damage in a Tropical Tree.” Proceedings of the National Academy of Sciences of the United States of America 100 (26): 15649–54. https://doi.org/10.1073/pnas.2533483100.

Aroca, Ángeles, David Gramaje, Josep Armengol, José García-Jiménez, and Rosa Raposo. (2010) “Evaluation of the Grapevine Nursery Propagation Process as a Source of *Phaeoacremonium* Spp. and *Phaeomoniella chlamydospora* and Occurrence of Trunk Disease Pathogens in Rootstock Mother Vines in Spain.” European Journal of Plant Pathology 126 (2): 165–74. https://doi.org/10.1007/s10658-009-9530-3.

Assante, Gemma, Dario Maffi, Marco Saracchi, Gandolfina Farena, Salvatore Moricca, and Alessandro Ragazzi. (2004) “Histological Studies on the Mycoparasitism of *Cladosporium tenuissimum* on Urediniospores of *Uromyees appendiculatus*” Mycological Research 108 (2): 170–82. https://doi.org/10.1017/S0953756203008852.

Auger, J., M. Esterio, I. Pérez, W. D. Gubler, and A. Eskalen. (2004) “First Report of *Phaeomoniella chlamydospora* on Vitis Vinifera and French American Hybrids in Chile.” Plant Disease 88 (11): 1285–1285. https://doi.org/10.1094/PDIS.2004.88.11.1285C.

Azevedo, João Lúcio, Walter Maccheroni, José Odair Pereira, and Welington Luiz De Araújo. (2000) “Endophytic Microorganisms: A Review on Insect Control and Recent Advances on Tropical Plants.” EJB Electronic Journal of Biotechnology 3 (1): 717–3458. http://www.ejb.org/content/vol3/issue1/full/4.

Badalyan, S.M, Innocenti, G., Garibyan, N.G. (2002) “Antagonistic Activity of Xylotrophic Mushrooms Against.” Phytopathologia Mediterranea 41 (April 2017): 220–25.

Barnett, H.L., and V. G Lilly. (1962) “A Destructive Mycoparasyte, *Gliocladium roseum*.” Mycological Society of America 54 (1): 72–77. http://www.jstor.org/stable/3756600.

Barnett, Horace.L., and Barry B. Hunter. (1955) “Illustrated Genera of Imperfect Fungi.” Transactions of the British Mycological Society. https://doi.org/10.1016/s0007-1536(55)80058-7.

Benítez, Tahía, Ana M. Rincón, M. Carmen Limón, and Antonio C. Codón. (2004) “Biocontrol Mechanisms of *Trichoderma* Strains.” International Microbiology 7 (4): 249–60. https://doi.org/1139-6709.

Bertsch, C., M. Ramírez-Suero, M. Magnin-Robert, P. Larignon, J. Chong, E. Abou-Mansour, A. Spagnolo, C. Clément, and F. Fontaine. (2013) “Grapevine Trunk Diseases: Complex and Still Poorly Understood.” Plant Pathology 62 (2): 243–65. https://doi.org/10.1111/j.1365-3059.2012.02674.x.

Besoain, X., C. Torres, G. A. Díaz, and B. A. Latorre. (2013) “First Report of *Neofusicoccum australe* Associated with Botryosphaeria Canker of Grapevine in Chile.” Plant Disease 97 (1): 143–143. https://doi.org/10.1094/pdis-07-12-0652-pdn.

Bester, W., P. W. Crous, and P. H. Fourie. (2007) “Evaluation of Fungicides as Potential Grapevine Pruning Wound Protectants against Botryosphaeria Species.” Australasian Plant Pathology 36 (1): 73–77. https://doi.org/10.1071/AP06086.

Billones-Baaijens, R., E. E. Jones, H. J. Ridgway, and M. V. Jaspers. (2013) “Virulence Affected by Assay Parameters during Grapevine Pathogenicity Studies with Botryosphaeriaceae Nursery Isolates.” Plant Pathology 62 (6): 1214–25. https://doi.org/10.1111/ppa.12051.

Borneman, J., and R. J. Hartin. (2000) “PCR Primers That Amplify Fungal RRNA Genes from Environmental Samples.” Applied and Environmental Microbiology 66 (10): 4356–60. https://doi.org/10.1128/AEM.66.10.4356-4360.2000.

Chambers, S. M. (1993) “Phytophthora root rot of Chestnut” [Doctorate Thesis] University of Adelaide, Australia.

Costadone, L., and W.D. Gubler. (2016) Biocontrol of Major Grapevine Diseases: Leading Research. Edited by Florence Mathieu Stephane Compant. Boston, MA: CABI.

Cota, L.V., L.A. Maffia, E.S.G. Mizubuti, and P.E.F. Macedo. (2009) “Biological Control by *Clonostachys rosea* as a Key Component in the Integrated Management of Strawberry Gray Mold.” Biological Control 50 (3): 222–30. https://doi.org/10.1016/j.biocontrol.2009.04.017.

Dennis, C., and J. Webster. (1971) “Antagonistic Properties of Species-Groups of *Trichoderma*.” Transactions of the British Mycological Society 57 (1): 25–IN3. https://doi.org/10.1016/s0007-1536(71)80077-3.

Díaz, G. A., and B. A. Latorre. (2014) “Infection Caused by *Phaeomoniella chlamydospora* Associated with Esca-like Symptoms in Grapevine in Chile.” Plant Disease 98 (3): 351–60. https://doi.org/10.1094/PDIS-12-12-1180-RE.

Díaz, G. A., D. Prehn, X. Besoain, E. R. Chávez, and B. A. Latorre. (2011) “*Neofusicoccum parvum* Associated with Grapevine Trunk Diseases in Chile.” Plant Disease 95 (8): 1032–1032. https://doi.org/10.1094/PDIS-03-11-0260.

Díaz, G. A., D. Prehn, and B. A. Latorre. (2011) “First Report of *Cryptovalsa ampelina* and *Eutypella leprosa* Associated with Grapevine Trunk Diseases in Chile.” Plant Disease 95 (4): 490–490. https://doi.org/10.1094/PDIS-12-10-0919.

Díaz, Gonzalo A., and Bernardo A. Latorre. (2013) “Efficacy of Paste and Liquid Fungicide Formulations to Protect Pruning Wounds against Pathogens Associated with Grapevine Trunk Diseases in Chile.” Crop Protection 46 (April): 106–12. https://doi.org/10.1016/j.cropro.2013.01.001.

Díaz, Gonzalo A, Jaime Auger, Ximena Besoain, Edmundo Bordeu, and Bernardo A. Latorre. (2013) “Prevalence and Pathogenicity of Fungi Associated with Grapevine Trunk Diseases in Chilean Vineyards.” Ciencia e Investigación Agraria 40 (2): 327–39. https://doi.org/10.4067/RCIA.V40I2.1101.

Fávaro, Léia Cecilia de Lima, Fernanda Luiza de Souza Sebastianes, and Welington Luiz Araújo. (2012) “*Epicoccum nigrum* P16, a Sugarcane Endophyte, Produces Antifungal Compounds and Induces Root Growth.” PloS One 7 (6): e36826. https://doi.org/10.1371/journal.pone.0036826.

Felzensztein, Christian. (2014) “The Chilean Wine Industry: New International Strategies for 2020.” Emerald Emerging Markets Case Studies 4 (2): 1–12. https://doi.org/10.1108/EEMCS-2014-2222.

Gao K., Gao, K, X Liu X., Liu, Z Kang Z., Kang, and K Mendgen K., Mendgen. (2005) “Mycoparasitism of *Rhizoctonia solani* by Endophytic *Chaetomium spirale* ND35: Ultrastructure and Cytochemistry of the Interaction.” Journal of Phytopathology 153 (5): 280–90. https://doi.org/10.1111/j.1439-0434.2005.00970.x.

Gramaje, D., and J. Armengol. (2011) “Importance and Impact of Fungal Trunk Pathogens in Young Vineyards.” Plant Disease 95 (9): 1040–55. https://doi.org/10.1074/jbc.M309555200.

Gramaje, David, Jose Ramon Urbez-Torres, and Mark R. Sosnowski. (2018) “Managing Grapevine Trunk Diseases with Respect to Etiology and Epidemiology: Current Strategies and Future Prospects.” Plant Disease 102 (1): 12–39. https://doi.org/10.1094/PDIS-04-17-0512-FE.

Guzmán, Bernardo Latorre. (2018) Compendio de las Enfermedades de las Plantas. 1st ed. Ediciones UC. http://www.jstor.org/stable/j.ctvkjb460.

Halleen, F., P. H. Fourie, and P. J. Lombard. (2010) “Protection of Grapevine Pruning Wounds against *Eutypa lata* by Biological and Chemical Methods.” South African Journal of Enology and Viticulture 31 (2): 125–32.

Hardoim, Pablo R., Leo S. van Overbeek, and Jan Dirk van Elsas. (2008) “Properties of Bacterial Endophytes and Their Proposed Role in Plant Growth.” Trends in Microbiology 16 (10): 463–71. https://doi.org/10.1016/j.tim.2008.07.008.

Hardoim, Pablo R., Leonard S. van Overbeek, Gabriele Berg, Anna Maria Pirttilä, Stéphane Compant, Andrea Campisano, Matthias Döring, and Angela Sessitsch. (2015) “The Hidden World within Plants: Ecological and Evolutionary Considerations for Defining Functioning of Microbial Endophytes.” Microbiology and Molecular Biology Reviews 79 (3): 293–320. https://doi.org/10.1128/MMBR.00050-14.

Howell, C. R. (2003) “Mechanisms Employed by *Trichoderma* Species in the Biological Control of Plant Diseases: The History and Evolution of Current Concepts.” Plant Disease 87 (1): 4–10. https://doi.org/10.1094/pdis.2003.87.1.4.

Hung, Phung Manh, Pongnak Wattanachai, Soytong Kasem, and Supattra Poeaim. (2015) “Efficacy of Chaetomium Species as Biological Control Agents against *Phytophthora nicotianae* Root Rot in Citrus.” Mycobiology 43 (3): 288. https://doi.org/10.5941/MYCO.2015.43.3.288.

Huu Phong, Nguyen, Wattanachai Pongnak, and Kasem Soytong. (2016) “Antifungal Activities of *Chaetomium* Spp. Against Fusarium Wilt of Tea.” Plant Protection Science 52 (1): 10–17. https://doi.org/10.17221/34/2015-PPS.

Ichi-Ishi, Akihiko, and Hirokazu Inoue. (2005) “Cloning, Nucleotide Sequence, and Expression of Tef-1, the Gene Encoding Translation Elongation Factor 1.ALPHA. (EF-1.ALPHA.) of *Neurospora crassa*.” The Japanese Journal of Genetics. https://doi.org/10.1266/jjg.70.273.

Inch, S, and J Gilbert. (2007) “Effect of Trichoderma Harzianum on Perithecial Production of *Gibberella zeae* on Wheat Straw.” Biocontrol Science and Technology 17 (6): 635–46. https://doi.org/10.1080/09583150701408865.

Iqbal, Mudassir, Mukesh Dubey, Kerstin McEwan, Uwe Menzel, Mikael Andersson Franko, Maria Viketoft, Dan Funck Jensen, and Magnus Karlsson. (2017) “Evaluation of *Clonostachys rosea* for Control of Plant-Parasitic Nematodes in Soil and in Roots of Carrot and Wheat” Phytopathology 108 (1): 52–59. https://doi.org/10.1094/phyto-03-17-0091-r.

Kaplan, Jonathan, Renaud Travadon, Monica Cooper, Vicken Hillis, Mark Lubell, and Kendra Baumgartner. (2016) “Identifying Economic Hurdles to Early Adoption of Preventative Practices: The Case of Trunk Diseases in California Winegrape Vineyards.” Wine Economics and Policy 5 (2): 127–41. https://doi.org/10.1016/j.wep.2016.11.001.

Karlsson, Magnus, Mikael Brandström Durling, Jaeyoung Choi, Chatchai Kosawang, Gerald Lackner, Georgios D. Tzelepis, Kristiina Nygren, et al. (2015) “Insights on the Evolution of Mycoparasitism from the Genome of *Clonostachys rosea*.” Genome Biology and Evolution 7 (2): 465–80. https://doi.org/10.1093/gbe/evu292.

Köhl, Jürgen, Christian Scheer, Imre J. Holb, Sylwester Masny, and Wilma Molhoek. (2015) “Toward an Integrated Use of Biological Control by *Cladosporium cladosporioides* H39 in Apple Scab (Venturia Inaequalis) Management.” Plant Disease 99 (4): 535–43. https://doi.org/10.1094/PDIS-08-14-0836-RE.

Latz, Meike A.C., Birgit Jensen, David B. Collinge, and Hans J.L. Jørgensen. (2018) “Endophytic Fungi as Biocontrol Agents: Elucidating Mechanisms in Disease Suppression.” Plant Ecology and Diversity 11 (5–6): 555–67. https://doi.org/10.1080/17550874.2018.1534146.

Lehtonen, Päivi T, Marjo Helander, Shahid A Siddiqui, Kirsi Lehto, and Kari Saikkonen. (2006) “Endophytic Fungus Decreases Plant Virus Infections in Meadow Ryegrass *(Lolium pratensef*.” Biology Letters 2 (4): 620–23. https://doi.org/10.1098/rsbl.2006.0499.

López-Fernández, Sebastián, Stéphane Compant, Urska Vrhovsek, Pier Luigi Bianchedi, Angela Sessitsch, Ilaria Pertot, and Andrea Campisano. 2016. “Grapevine Colonization by Endophytic Bacteria Shifts Secondary Metabolism and Suggests Activation of Defense Pathways.” Plant and Soil 405 (1–2): 155–75. https://doi.org/10.1007/s11104-015-2631-1.

Massonnet, Mélanie, Rosa Figueroa-Balderas, Erin R.A. Galarneau, Shiho Miki, Daniel P. Lawrence, Qiang Sun, Christopher M. Wallis, Kendra Baumgartner, and Dario Cantu. (2017) “*Neofusicoccum parvum* Colonization of the Grapevine Woody Stem Triggers Asynchronous Host Responses at the Site of Infection and in the Leaves.” Frontiers in Plant Science 8(June). https://doi.org/10.3389/fpls.2017.01117.

Mondello, Vincenzo, Aurélie Songy, Enrico Battiston, Catia Pinto, Cindy Coppin, Patricia Trotel-Aziz, Christophe Clément, Laura Mugnai, and Florence Fontaine. (2018) “Grapevine Trunk Diseases: A Review of Fifteen Years of Trials for Their Control with Chemicals and Biocontrol Agents.” Plant Disease 102 (7): 1189–1217. https://doi.org/10.1094/PDIS-08-17-1181-FE.

Moraga-Suazo, P, Sanfuentes E, and Le-Feuvre R. (2016) “Induced Systemic Resistance Triggered by *Clonostachys rosea* Against *Fusarium circinatum* in *Pinus radiata*.” Forest Research: Open Access 5 (2): 1–4. https://doi.org/10.4172/2168-9776.1000174.

Morandi, M A B, L A Maffia, and J C Sutton. (2001) “Development of *Clonostachys rosea* and Interactions with *Botrytis cinerea* in Rose Leaves and Residues.” Phytoparasitica 29 (2): 103–13.

Mousa, Walaa Kamel, and Manish N Raizada. (2013) “The Diversity of Anti-Microbial Secondary Metabolites Produced by Fungal Endophytes: An Interdisciplinary Perspective.” Frontiers in Microbiology 4 (March): 65. https://doi.org/10.3389/fmicb.2013.00065.

Munkvold, G. P., J. A. Duthie, and J. J. Marois. (1994) “Reductions in Yield and Vegetative Growth of Grapevines Due to Eutypa Dieback.” The American Phytopathological Society 84 (2): 186–92. https://doi.org/10.1094/Phyto-84-186.

Nilsson, Rolf Henrik, Karl Henrik Larsson, Andy F.S. Taylor, Johan Bengtsson-Palme, Thomas S. Jeppesen, Dmitry Schigel, Peter Kennedy. (2019) “The UNITE Database for Molecular Identification of Fungi: Handling Dark Taxa and Parallel Taxonomic Classifications.” Nucleic Acids Research 47 (D1): D259–64. https://doi.org/10.1093/nar/gky1022.

Pancher, Michael, Marco Ceol, Paola Elisa Corneo, Claudia Maria Oliveira Longa, Sohail Yousaf, Ilaria Pertot, and Andrea Campisano. (2012) “Fungal Endophytic Communities in Grapevines (Vitis Vinifera L.) Respond to Crop Management.” Applied and Environmental Microbiology 78 (12): 4308–17. https://doi.org/10.1128/AEM.07655-11.

Petrini, Orlando. (1991) “Fungal Endophytes of Tree Leaves,” 179–97. https://doi.org/10.1007/978-1-4612-3168-4_9.

Petzoldt, C. H. 1981. “Eutypa Dieback of Grapevine: Seasonal Differences in Infection and Duration of Susceptibility of Pruning Wounds.” Phytopathology. https://doi.org/10.1094/phyto-71-540.

Pieterse, Corné M. J., Zamioudis, Christos, Berendsen, Roeland L., Weller, David M., Van Wees, Saskia C. M., Bakker, Peter A. H. M. (2014) “Induced Systemic Resistance by Beneficial Microbes.” Annual Review of Phytopathology 52 (1): 347–375. https://doi.org/10.1146/anurev-phyto-082712-102340

Pizarro, María José. (2018) “Boletín Del Vino: Producción, Precios y Comercio Exterior. Avance a Octubre de 2018.” ODEPA, Ministerio de Agricultura, Gobierno de Chile.

Rodriguez, M. A., G. Cabrera, F. C. Gozzo, M. N. Eberlin, and A. Godeas. (2011) “*Clonostachys rosea* BAFC3874 as a Sclerotinia Sclerotiorum Antagonist: Mechanisms Involved and Potential as a Biocontrol Agent.” Journal of Applied Microbiology 110 (5): 1177–86. https://doi.org/10.1111/j.1365-2672.2011.04970.x.

Rolshausen, P. E., and W. D. Gubler. (2005) “Use of Boron for the Control of Eutypa Dieback of Grapevines.” Plant Disease 89 (7): 734–38. https://doi.org/10.1094/PD-89-0734.

Rolshausen, Philippe E., Jose Ramon Urbez-Torres, Suzanne Rooney-Latham, Akif Eskalen, Rhonda J. Smith, and Walter Douglas Gubler. (2010) “Evaluation of Pruning Wound Susceptibility and Protection against Fungi Associated with Grapevine Trunk Diseases.” American Journal of Enology and Viticulture 61 (1): 113–19.

Schindelin, Johannes, Ignacio Arganda-Carreras, Erwin Frise, Verena Kaynig, Mark Longair, Tobias Pietzsch, Stephan Preibisch. (2012) “Fiji: An Open-Source Platform for Biological-Image Analysis.” Nature Methods 9 (June): 676. https://doi.org/10.1038/nmeth.2019.

Schöneberg, A., T. Musa, R. T. Voegele, and S. Vogelgsang. (2015a) “The Potential of Antagonistic Fungi for Control of Fusarium Graminearum and Fusarium Crookwellense Varies Depending on the Experimental Approach.” Journal of Applied Microbiology 118 (5): 1165–79. https://doi.org/10.1111/jam.12775.

Schöneberg, A., T. Musa, R. T. Voegele, and S. Vogelgsang. (2015b) “The Potential of Antagonistic Fungi for Control of Fusarium Graminearum and Fusarium Crookwellense Varies Depending on the Experimental Approach.” Journal of Applied Microbiology 118 (5): 1165–79. https://doi.org/10.1111/jam.12775.

Schulz, Barbara, and Christine Boyle. (2005) “The Endophytic Continuum.” Mycological Research 109 (6): 661–86. https://doi.org/10.1017/S095375620500273X.

Sharmini, John, Scott, Eileen S., Wicks, Trevor J. and Hunt, John S. (2004) “Interactions between *Eutypa lata* and *Trichoderma harzianum*.” Phytopathologia Mediterranea 43 (1): 95–104.

Solano Castillo I, Tulio F, Marcia L Castillo Ávila I, José V Medina Medina I, and Elio M Del Pozo Núñez. (2014) “Efectividad de Hongos Nematófagos Sobre *Meloidogyne incognita* (Kofoid y White) Chitwood En Tomate En Condiciones de Campo, Loja, Ecuador Effectiveness of Nematophagous Fungi on *Meloidogyne incognita* (Kofoid and White) Chitwood on Tomato in Field Conditions in Loja, Ecuador.” Rev. Protección Veg 29 (3): 192–96. http://scielo.sld.cu/pdf/rpv/v29n3/rpv05314.pdf.

Sosnowski, Mark R., and Dion C. Mundy. (2018) “Pruning Wound Protection Strategies for Simultaneous Control of Eutypa and Botryosphaeria Dieback in New Zealand.” Plant Disease 103 (3): 1–26. https://doi.org/https://doi.org/10.1094/PDIS-05-18-0728-RE.

Sosnowski, Mark R., and Dion C. Mundy. (2019) “Pruning Wound Protection Strategies for Simultaneous Control of Eutypa and Botryosphaeria Dieback in New Zealand.” Plant Disease 103 (3): 519–25. https://doi.org/10.1094/PDIS-05-18-0728-RE.

Surico, Giuseppe, Laura Mugnai, and Guido Marchi. (2006) “Older and More Recent Observations on Esca: A Critical Overview.” Phytopathologia Mediterranea 45 (SUPPL. 1): 68–86. https://doi.org/10.14601/PHYTOPATHOL_MEDITERR-1847.

Tan, Yong, Yinshan Cui, Haoyu Li, Anxiu Kuang, Xiaoran Li, Yunlin Wei, and Xiuling Ji. (2017) “Rhizospheric Soil and Root Endogenous Fungal Diversity and Composition in Response to Continuous Panax Notoginseng Cropping Practices.” Microbiological Research 194: 10–19. https://doi.org/10.1016/j.micres.2016.09.009.

Travadon, Renaud, Pascal Lecomte, Barka Diarra, Daniel P. Lawrence, David Renault, Hernán Ojeda, Patrice Rey, and Kendra Baumgartner. (2016) “Grapevine Pruning Systems and Cultivars Influence the Diversity of Wood-Colonizing Fungi.” Fungal Ecology 24: 82–93. https://doi.org/10.1016/j.funeco.2016.09.003.

USDA Foreign Agricultural Center. (2019) “Grapes, Fresh Table: Production, Supply and Distribution.” https://apps.fas.usda.gov/psdonline/reporthandler.ashx?reportId=2419&templateId=3&format=html&fileName=Grapes [Accessed March 11, 2019].

Wagschal, Isabelle, Eliane Abou-Mansour, Anne-Noelle Petit, Christophe Clement, and Florence Fontaine. (2008) “Wood Diseases of Grapevine: A Review on Eutypa Dieback and Esca.” Research Signpost 37 (2): 661–86.

Waite, Helen, and Lucie Morton. (2007) “Hot Water Treatment, Trunk Diseases and Other Critical Factors in the Production of High-Quality Grapevine Planting Material.” Phytopathologia Mediterranea 46 (1): 5–17. https://doi.org/10.14601/phytopathol_mediterr-1857.

Weber, Edward A, Florent P Trouillas, and W Douglas Gubler. (2007) “Double Pruning of Grapevines: A Cultural Practice to Reduce Infections by *Eutypa lata*” American Journal of Enology and Viticulture 58 (1): 61 LP–66. http://www.ajevonline.org/content/58/1/61.abstract.

White, TJ., T. Bruns, SW. Lee, and JW. Taylor. (1990) “Amplification and Direct Sequencing of Fungal Ribosomal RNA Genes for Phylogenetics.” In PCR Protocols: A Guide to Methods and Applications, edited by MA Innis, DH Gelfand, JJ Sninsky, and TJ White, 315–22. New York, NY.: Academic Press Inc.

Wicaksono, Wisnu Adi, E. Eirian Jones, Jana Monk, and Hayley J. Ridgway. (2017) “Using Bacterial Endophytes from a New Zealand Native Medicinal Plant for Control of Grapevine Trunk Diseases.” Biological Control 114 (August): 65–72. https://doi.org/10.1016/j.biocontrol.2017.08.003.

Wightwick, Adam, Robert Walters, Graeme Allinson, Suzanne Reichman, and Neal Menzies. (2010) “Environmental Risks of Fungicides Used in Horticultural Production Systems.” Fungicides. https://doi.org/10.5772/13032.

Zabalgogeazcoa, I. (2008) “Fungal Endophytes and Their Interaction with Plant Pathogens.” Spanish Journal of Agricultural Research 6: 138–46. https://doi.org/10.5424/sjar/200806S1-382.

