## Supplemental Table 1 for "Biocontrol potential of grapevine endophytes against grapevine trunk pathogens"

### *Supplementary Material*

**Supplementary Table 1**: Isolate's first screening against *Diplodia seriata*

| Disease Control | Origin | Sample | Isolate | *D. seriata* growth inhibition (%) |
|---|---|---|---|---|
| None | Sprouts | 1 | 1b | 17,43 |
| None | Sprouts | 1 | 3d | 30,43 |
| None | Sprouts | 1 | 1d | 32,33 |
| None | Sprouts | 1 | 2b | 35,94 |
| Conventional | Sprouts | 1 | 2v | 37,19 |
| Conventional | Sprouts | 1 | 2 | 38,25 |
| None | Root | 1 | 5 | 19,34 |
| None | Root | 1 | 11 | 25,20 |
| None | Root | 1 | 12 | 25,43 |
| None | Root | 1 | 6 | 27,22 |
| None | Root | 1 | 13 | 28,47 |
| None | Root | 1 | 9 | 36,01 |
| None | Root | 1 | 16 | 36,09 |
| None | Root | 1 | 18 | 64,91 |
| Conventional | Root | 1 | 1 | 51,2 |
| Conventional | Root | 1 | 2 | no |
| None | Rizospheric soil | 1 | 15 | 14,82 |
| None | Rizospheric soil | 1 | 21 | 18,64 |
| None | Rizospheric soil | 1 | 32 | 19,37 |

| | | | | |
|---|---|---|---|---|
| None | Rizospheric soil | 1 | 30 | 20,77 |
| None | Rizospheric soil | 1 | 11 | 21,76 |
| None | Rizospheric soil | 1 | 19 | 27,90 |
| None | Rizospheric soil | 1 | 29 | 34,50 |
| None | Rizospheric soil | 1 | 31 | 37,50 |
| None | Rizospheric soil | 1 | 28 | 39,00 |
| None | Rizospheric soil | 1 | 9 | 39,25 |
| None | Rizospheric soil | 1 | 33 | 39,85 |
| None | Rizospheric soil | 1 | 20 | 40,40 |
| None | Rizospheric soil | 1 | 16 | 44,17 |
| None | Rizospheric soil | 1 | 25 | 51,66 |
| None | Rizospheric soil | 1 | 6 | 58,04 |
| None | Rizospheric soil | 1 | 23 | 58,15 |
| None | Rizospheric soil | 1 | 14 | 77,83 |
| None | Rizospheric soil | 1 | 4 | 84,86 |
| None | Rizospheric soil | 1 | 3 | 84,98 |
| None | Sprouts | 2 | 1d | 36,65 |
| Conventional | Sprouts | 2 | 1 | 19,91 |
| None | Root | 2 | 20 | 3,78 |
| None | Root | 2 | 3 | 12,84 |
| None | Root | 2 | 27 | 14,56 |
| None | Root | 2 | 26v | 15,05 |
| None | Root | 2 | 4 | 18,54 |

| | | | | |
|---|---|---|---|---|
| None | Root | 2 | 15 | 19,40 |
| None | Root | 2 | 11 | 27,60 |
| None | Root | 2 | 28 | 29,84 |
| None | Root | 2 | 7 | 35,44 |
| None | Root | 2 | 17 | 36,04 |
| None | Root | 2 | 11 | 36,96 |
| None | Root | 2 | 13 | 38,25 |
| None | Root | 2 | 6 | 40,78 |
| None | Root | 2 | 1 | 41,54 |
| None | Root | 2 | 8 | 41,58 |
| None | Root | 2 | 21 | 42,93 |
| None | Root | 2 | 24 | 43,93 |
| None | Root | 2 | 26 | 44,85 |
| None | Root | 2 | 13 | 47,11 |
| None | Root | 2 | 5 | 54,84 |
| None | Root | 2 | 10 | 59,61 |
| None | Root | 2 | 12 | 66,38 |
| None | Root | 2 | 23 | 78,51 |
| Conventional | Root | 2 | 2 | 16,29 |
| Conventional | Root | 2 | 6 | 31,03 |
| Conventional | Root | 2 | 7 | 33,17 |
| Conventional | Root | 2 | 8v | 38,4 |
| Conventional | Root | 2 | 1 | 40,11 |

| | | | | |
|---|---|---|---|---|
| Conventional | Root | 2 | 8 | 44,9 |
| Conventional | Root | 2 | 5 | 61,22 |
| None | Rizospheric soil | 2 | 23 | 13,83 |
| None | Rizospheric soil | 2 | 2 | 17,94 |
| None | Rizospheric soil | 2 | 11 | 21,16 |
| None | Rizospheric soil | 2 | 7 | 21,33 |
| None | Rizospheric soil | 2 | 3 | 22,05 |
| None | Rizospheric soil | 2 | 28 | 22,21 |
| None | Rizospheric soil | 2 | 15 | 22,71 |
| None | Rizospheric soil | 2 | 5 | 24,95 |
| None | Rizospheric soil | 2 | 6 | 25,61 |
| None | Rizospheric soil | 2 | 4 | 29,97 |
| None | Rizospheric soil | 2 | 12 | 30,01 |
| None | Rizospheric soil | 2 | 19 | 37,61 |
| None | Rizospheric soil | 2 | 1 | 38,77 |
| None | Rizospheric soil | 2 | 25 | 40,15 |
| None | Rizospheric soil | 2 | 17 | 43,18 |
| None | Rizospheric soil | 2 | 10 | 61,23 |
| None | Rizospheric soil | 2 | 13 | 70,31 |
| None | Rizospheric soil | 2 | 20 | 79,69 |
| Conventional | Rizospheric soil | 2 | 1v | 35,183 |
| Conventional | Rizospheric soil | 2 | 1 | 38,02 |
| Conventional | Sprouts | 3 | 1 | 42,46 |

| | | | | |
|---|---|---|---|---|
| Conventional | Root | 3 | 4 | 25,95 |
| Conventional | Root | 3 | 2 | no |
| Conventional | Rizospheric soil | 3 | 2 | 27,43 |
| None | Sprouts | 4 | 2d | 22,67 |
| None | Sprouts | 4 | 3d | 28,434 |
| Conventional | Sprouts | 4 | 1 | 18,63 |
| Conventional | Sprouts | 5 | 1 | 34,06 |
| Conventional | Sprouts | 5 | 1 | 51,11 |
| None | Root | 5 | 2 | 39,29 |
| None | Root | 5 | 2 | 93,02 |
| Conventional | Root | 5 | 3 | 21,11 |
| Conventional | Root | 5 | 4 | 23,95 |
| None | Rizospheric soil | 5 | 6 | 7,89 |
| None | Rizospheric soil | 5 | 15 | 9,66 |
| None | Rizospheric soil | 5 | 5 | 12,27 |
| None | Rizospheric soil | 5 | 19 | 15,96 |
| None | Rizospheric soil | 5 | 3 | 16,27 |
| None | Rizospheric soil | 5 | 11 | 17,26 |
| None | Rizospheric soil | 5 | 18 | 28,79 |
| None | Rizospheric soil | 5 | 14 | 40,93 |
| None | Rizospheric soil | 5 | 1 | 42,92 |
| None | Rizospheric soil | 5 | 7 | 68,72 |
| None | Rizospheric soil | 5 | 21 | 97,56 |

| | | | | |
|---|---|---|---|---|
| None | Sprouts | 6 | 2d | 27,57 |
| None | Sprouts | 6 | 1a | 39,42 |
| None | Root | 6 | 3 | 11,71 |
| None | Root | 6 | 2 | 13,26 |
| None | Root | 6 | 5 | 15,47 |
| None | Root | 6 | 1 | 35,14 |
| Conventional | Root | 6 | 1 | 39,65 |
| Conventional | Root | 6 | 2 | 49,71 |
| None | Rizospheric soil | 6 | 6 | 5,66 |
| None | Rizospheric soil | 6 | 29 | 8,11 |
| None | Rizospheric soil | 6 | 23 | 10,27 |
| None | Rizospheric soil | 6 | 11 | 13,41 |
| None | Rizospheric soil | 6 | 31 | 14,56 |
| None | Rizospheric soil | 6 | 3 | 15,76 |
| None | Rizospheric soil | 6 | 26 | 16,02 |
| None | Rizospheric soil | 6 | 22 | 18,74 |
| None | Rizospheric soil | 6 | 16 | 22,93 |
| None | Rizospheric soil | 6 | 2 | 24,23 |
| None | Rizospheric soil | 6 | 15 | 28,70 |
| None | Rizospheric soil | 6 | 33 | 30,15 |
| None | Rizospheric soil | 6 | 19 | 32,20 |
| None | Rizospheric soil | 6 | 8 | 32,29 |
| None | Rizospheric soil | 6 | 34 | 36,45 |

| | | | | |
|---|---|---|---|---|
| None | Rizospheric soil | 6 | 32 | 36,57 |
| None | Rizospheric soil | 6 | 21 | 49,11 |
| None | Rizospheric soil | 6 | 27 | 51,97 |
| None | Rizospheric soil | 6 | 1 | 57,28 |
| None | Rizospheric soil | 6 | 9 | 58,72 |
| None | Rizospheric soil | 6 | 4 | 59,40 |
| None | Rizospheric soil | 6 | 12 | 62,38 |
| None | Rizospheric soil | 6 | 31v | 75,01 |
| None | Rizospheric soil | 6 | 28 | 92,82 |
| None | Rizospheric soil | 6 | 20 | 97,96 |
| Conventional | Rizospheric soil | 6 | 2 | 7,43 |
| None | Root | 7 | 11 | 3,45 |
| None | Root | 7 | 10 | 14,51 |
| None | Root | 7 | 12 | 18,71 |
| None | Root | 7 | 5 | 27,82 |
| None | Root | 7 | 9 | 29,14 |
| None | Root | 7 | 13 | 39,64 |
| None | Root | 7 | 1 | 58,39 |
| None | Root | 7 | 5 | 61,69 |
| Conventional | Root | 7 | 2 | 17,2 |
| None | Rizospheric soil | 7 | 3 | 12,43 |
| None | Rizospheric soil | 7 | 5 | 28,08 |
| None | Rizospheric soil | 7 | 1 | 32,13 |

| | | | | |
|---|---|---|---|---|
| None | Rizospheric soil | 7 | 6 | 33,80 |
| None | Rizospheric soil | 7 | 8 | 44,42 |
| None | Rizospheric soil | 7 | 7 | 44,83 |
| None | Rizospheric soil | 7 | 23 | 45,29 |
| None | Rizospheric soil | 7 | 9 | 48,00 |
| Conventional | Rizospheric soil | 7 | 1 | 44,36 |
| Conventional | Root | 8 | 2 | 31,46 |
| Conventional | Root | 8 | 1 | no |
| Conventional | Rizospheric soil | 8 | 1 | 33,18 |
| Conventional | Rizospheric soil | 8 | 2 | 78,91 |
| Conventional | Root | 9 | 2 | 32,9 |
| Conventional | Root | 10 | 1 | 38,98 |
| Organic | Sprouts | 11 | 1a | 11,38 |
| Organic | Sprouts | 11 | 2a | 33,12 |
| Organic | Sprouts | 11 | 3 | 41,7 |
| Organic | Rizospheric soil | 11 | 4 | 28,543 |
| Organic | Rizospheric soil | 11 | 7 | 37,56 |
| Organic | Rizospheric soil | 11 | 3 | 42,02 |
| Organic | Rizospheric soil | 11 | 1 | 42,99 |
| Organic | Rizospheric soil | 11 | 6 | 51,92 |
| Organic | Root | 13 | 1 | 21,94 |
| Organic | Root | 13 | 5 | 29,16 |
| Organic | Root | 13 | 2 | 29,3 |

| | | | | |
|---|---|---|---|---|
| Organic | Root | 13 | 3 | 40,69 |
| Organic | Sprouts | 14 | 1a | 19,28 |
| Organic | Root | 15 | 2 | 31,22 |
| Organic | Root | 15 | 1 | 38,76 |
| Organic | Sprouts | 16 | 1 | 37,18 |
| Organic | Root | 16 | 2 | 21,03 |
| Organic | Root | 16 | 3v | 36,24 |
| Organic | Root | 16 | 3 | 61,22 |
| Organic | Root | 16 | 1 | 66,83 |
| Organic | Rizospheric soil | 16 | 1v | 19,32 |
| Organic | Rizospheric soil | 16 | 1 | 33,01 |
| Organic | Sprouts | 20 | 2 | 4,01 |
| Organic | Sprouts | 20 | 1 | 28,47 |
| Organic | Root | 20 | 1 | 18,03 |
| Organic | Root | 20 | 1v | 25,29 |
| Organic | Root | 20 | 2 | 51,64 |
| Organic | Root | 20 | 3 | 67,15 |
| Organic | Rizospheric soil | 20 | 15 | 16,3 |
| Organic | Rizospheric soil | 20 | 14 | 23,22 |
| Organic | Rizospheric soil | 20 | 3 | 25,38 |
| Organic | Rizospheric soil | 20 | 1 | 38,01 |
| Organic | Rizospheric soil | 20 | 13 | 40,32 |
| Organic | Rizospheric soil | 20 | 4 | 41,52 |

| | | | | |
|---|---|---|---|---|
| Organic | Rizospheric soil | 20 | 6 | 46,98 |
| Organic | Rizospheric soil | 20 | 8 | 48,9 |
| Organic | Sprouts | 22 | 2a | 9,45 |
| Organic | Sprouts | 22 | 2 | 22,91 |
| Organic | Root | 22 | 2 | 43,37 |
| Organic | Rizospheric soil | 22 | 3 | 9,7 |
| Organic | Rizospheric soil | 22 | 1 | 44,99 |
| Organic | Root | 25 | 2 | 27,30 |
| Organic | Root | 25 | 2v | 29,36 |
| Organic | Root | 25 | 3 | 42,93 |
| Organic | Rizospheric soil | 25 | 3 | 30,05 |
| Organic | Rizospheric soil | 25 | 1 | 33,04 |
| Organic | Rizospheric soil | 25 | 5 | 33,65 |
| Organic | Sprouts | 27 | 1a | 43,02 |
| Organic | Root | 27 | 3 | 7,12 |
| Organic | Root | 27 | 2 | 8,23 |
| Organic | Root | 27 | 1 | 31,02 |
| Organic | Rizospheric soil | 27 | 3 | 13,43 |
| Organic | Rizospheric soil | 27 | 2 | 29,35 |
| Organic | Rizospheric soil | 27 | 3 | 45,23 |
| Organic | Rizospheric soil | 27 | 1 | 48,20 |
| Organic | Root | 29 | 1 | 17,54 |
| Organic | Root | 30 | 3 | 38,9 |

| | | | | |
|---|---|---|---|---|
| Organic | Root | 30 | 2 | 40,78 |
| Organic | Rizospheric soil | 30 | 5 | 7,91 |
| Organic | Rizospheric soil | 30 | 4v | 30,17 |
| Organic | Rizospheric soil | 30 | 1 | 39,43 |
| Organic | Rizospheric soil | 30 | 2v | 42,93 |
| Organic | Rizospheric soil | 30 | 2 | 44,58 |
| Organic | Rizospheric soil | 30 | 4 | 55,21 |
| Conventional | Root | 31 | 1 | 19,45 |
| Conventional | Root | 31 | 3 | 27,19 |
| Conventional | Root | 31 | 6 | 31,28 |
| Conventional | Root | 31 | 5 | 47,19 |
| Conventional | Root | 31 | 7 | 88,1 |
| Conventional | Rizospheric soil | 31 | 10 | 15,87 |
| Conventional | Rizospheric soil | 31 | 7 | 22,48 |
| Conventional | Rizospheric soil | 31 | 9 | 23,95 |
| Conventional | Rizospheric soil | 31 | 10v | 25,37 |
| Conventional | Rizospheric soil | 31 | 1 | 27,38 |
| Conventional | Rizospheric soil | 31 | 5 | 38,44 |
| Conventional | Rizospheric soil | 31 | 14 | 41,65 |
| Conventional | Rizospheric soil | 31 | 3 | 42,2978 |
| Conventional | Rizospheric soil | 31 | 7 | 45,2 |
| Conventional | Rizospheric soil | 31 | 12 | 48,46 |
| Conventional | Rizospheric soil | 31 | 6 | 55,01 |

| | | | | |
|---|---|---|---|---|
| Conventional | Rizospheric soil | 31 | 13 | 69,15 |
| Conventional | Sprouts | 32 | 2 | 35,19 |
| Conventional | Sprouts | 32 | 1 | 38,19 |
| Conventional | Root | 32 | 3 | 29,37 |
| Conventional | Root | 32 | 1 | 39,1 |
| Conventional | Rizospheric soil | 32 | 4 | 41,11 |
| Conventional | Rizospheric soil | 32 | 1 | 47,73 |
| Conventional | Root | 33 | 7 | 15,02 |
| Conventional | Root | 33 | 3 | 19,2 |
| Conventional | Root | 33 | 3 | 27,39 |
| Conventional | Root | 33 | 11v | 32,53 |
| Conventional | Root | 33 | 2 | 40,2 |
| Conventional | Root | 33 | 2v | 47,19 |
| Conventional | Root | 33 | 12 | 47,76 |
| Conventional | Root | 33 | 9 | 55,8 |
| Conventional | Root | 33 | 5 | 59,67 |
| Conventional | Root | 33 | 11 | no |
| Conventional | Rizospheric soil | 33 | 15 | 36,50 |
| Conventional | Rizospheric soil | 33 | 5 | 37,28 |
| Conventional | Rizospheric soil | 33 | 12 | 37,30 |
| Conventional | Rizospheric soil | 33 | 14 | 38,22 |
| Conventional | Rizospheric soil | 33 | 6 | 41,99 |
| Conventional | Rizospheric soil | 33 | 4 | 42,07 |

| | | | | |
|---|---|---|---|---|
| Conventional | Rizospheric soil | 33 | 13 | 44,61 |
| Conventional | Rizospheric soil | 33 | 7 | 48,26 |
| Conventional | Rizospheric soil | 33 | 18 | 48,29 |
| Conventional | Rizospheric soil | 33 | 4 | 48,56 |
| Conventional | Rizospheric soil | 33 | 3 | 51,03 |
| Conventional | Sprouts | 34 | 2c | 19,35 |
| Conventional | Sprouts | 34 | 1d | 24,88 |
| Conventional | Sprouts | 34 | 1a | 32,94 |
| Conventional | Sprouts | 34 | 3 | 41,82 |
| Conventional | Sprouts | 34 | 3 | no |
| Conventional | Root | 34 | 3 | 4,03 |
| Conventional | Root | 34 | 7 | 12,39 |
| Conventional | Root | 34 | 2 | 21,06 |
| Conventional | Root | 34 | 3 | 28,99 |
| Conventional | Root | 34 | 1 | 30,57 |
| Conventional | Root | 34 | 6 | 32,01 |
| Conventional | Root | 34 | 2v | 37,29 |
| Conventional | Root | 34 | 10 | 83,02 |
| Conventional | Rizospheric soil | 34 | 1 | 9,01 |
| Conventional | Rizospheric soil | 34 | 22 | 11,83 |
| Conventional | Rizospheric soil | 34 | 2 | 15,29 |
| Conventional | Rizospheric soil | 34 | 10 | 22,19 |
| Conventional | Rizospheric soil | 34 | 9 | 33,29 |

| | | | | |
|---|---|---|---|---|
| Conventional | Rizospheric soil | 34 | 4 | 33,85 |
| Conventional | Rizospheric soil | 34 | 20 | 36,45 |
| Conventional | Rizospheric soil | 34 | 7 | 38,28 |
| Conventional | Rizospheric soil | 34 | 24 | 40,11 |
| Conventional | Rizospheric soil | 34 | 16 | 41,01 |
| Conventional | Rizospheric soil | 34 | 19 | 41,22 |
| Conventional | Rizospheric soil | 34 | 15 | 44,73 |
| Conventional | Rizospheric soil | 34 | 23 | 48,12 |
| Conventional | Rizospheric soil | 34 | 8 | 52,06 |
| Conventional | Rizospheric soil | 34 | 6 | 57,02 |
| Conventional | Rizospheric soil | 35 | 10 | 11,03 |
| Conventional | Rizospheric soil | 35 | 4 | 17,29 |
| Conventional | Rizospheric soil | 35 | 5 | 28,45 |
| Conventional | Rizospheric soil | 35 | 14 | 36,187 |
| Conventional | Rizospheric soil | 35 | 11 | 37,29 |
| Conventional | Rizospheric soil | 35 | 13 | 47,25 |
| Conventional | Rizospheric soil | 35 | 9 | 62,94 |
| Conventional | Rizospheric soil | 35 | 7 | 71,92 |
| Conventional | Sprouts | 36 | 2a | 37,25 |
| Organic | Sprouts | 36 | 1 | 46,29 |
| Organic | Rizospheric soil | 36 | 1 | 9,17 |
| Organic | Rizospheric soil | 36 | 1 | 27,1 |
| Organic | Root | 37 | 1 | 11,24 |

| | | | | |
|---|---|---|---|---|
| Organic | Root | 37 | 2 | 18,73 |
| Organic | Root | 37 | 5 | 37,92 |
| Organic | Sprouts | 38 | 1c | 26,1 |
| Organic | Sprouts | 38 | 2 | 29,93 |
| Organic | Root | 38 | 1 | 19,22 |
| Organic | Root | 38 | 5 | 26,382 |
| Organic | Root | 38 | 4 | 31,5 |
| Organic | Root | 38 | 6v | 44,92 |
| Organic | Root | 38 | 4v | 46,1 |
| Organic | Root | 38 | 7 | 59,16 |
| Organic | Root | 38 | 6 | 65,92 |
| Organic | Rizospheric soil | 38 | 2 | 33,36 |
| Organic | Rizospheric soil | 38 | 4 | 40,81 |
| Organic | Rizospheric soil | 38 | 1 | 42,93 |
| Organic | Rizospheric soil | 38 | 7 | 47,02 |
| Organic | Sprouts | 39 | 3d | 33,67 |
| Organic | Root | 39 | 4 | 26,01 |
| Organic | Root | 39 | 3v | 26,36 |
| Organic | Root | 39 | 2 | 40,7 |
| Organic | Root | 39 | 1 | 52,05 |
| Organic | Root | 39 | 3 | 67,01 |
| Organic | Rizospheric soil | 39 | 1 | 71,28 |
| Organic | Sprouts | 40 | 1 | 42,3 |

| | | | | |
|---|---|---|---|---|
| Organic | Sprouts | 40 | 2 | 45,01 |
| Organic | Root | 40 | 7 | 49,62 |
| Organic | Root | 40 | 2 | 52 |
| Organic | Rizospheric soil | 40 | 1 | 29,345 |
| Organic | Rizospheric soil | 40 | 2 | 31,88 |
| Organic | Root | 41 | 3 | 21,03 |
| Organic | Root | 41 | 4 | 27,4876 |
| Organic | Root | 41 | 2 | 39,22 |
| Organic | Root | 41 | 1 | 55,81 |
| Organic | Rizospheric soil | 41 | 10 | 22,56 |
| Organic | Rizospheric soil | 41 | 4 | 28,43 |
| Organic | Rizospheric soil | 41 | 8 | 39,27 |
| Organic | Rizospheric soil | 41 | 7 | 39,59 |
| Organic | Rizospheric soil | 41 | 9 | 44,17 |
| Organic | Rizospheric soil | 41 | 11 | 53,95 |
| None | Root | 3 y 4 | 2 | 27,55 |
| None | Root | 3 y 4 | 3 | 36,21 |
| None | Root | 3 y 4 | 1 | 72,68 |
| None | Root | 3 y 4 | 7 | 90,43 |
| None | Rizospheric soil | 3 y 4 | 30 | 3,45 |
| None | Rizospheric soil | 3 y 4 | 20 | 7,73 |
| None | Rizospheric soil | 3 y 4 | 24 | 20,71 |
| None | Rizospheric soil | 3 y 4 | 23v | 22,35 |

| | | | | |
|---|---|---|---|---|
| None | Rizospheric soil | 3 y 4 | 12 | 22,79 |
| None | Rizospheric soil | 3 y 4 | 25 | 22,92 |
| None | Rizospheric soil | 3 y 4 | 4 | 24,59 |
| None | Rizospheric soil | 3 y 4 | 14 | 25,58 |
| None | Rizospheric soil | 3 y 4 | 13 | 26,68 |
| None | Rizospheric soil | 3 y 4 | 29 | 28,66 |
| None | Rizospheric soil | 3 y 4 | 28 | 31,84 |
| None | Rizospheric soil | 3 y 4 | 8 | 32,70 |
| None | Rizospheric soil | 3 y 4 | 5 | 32,71 |
| None | Rizospheric soil | 3 y 4 | 18 | 33,19 |
| None | Rizospheric soil | 3 y 4 | 23 | 34,73 |
| None | Rizospheric soil | 3 y 4 | 27 | 36,91 |
| None | Rizospheric soil | 3 y 4 | 22 | 37,32 |
| None | Rizospheric soil | 3 y 4 | 21v | 37,67 |
| None | Rizospheric soil | 3 y 4 | 11 | 39,91 |
| None | Rizospheric soil | 3 y 4 | 10 | 40,89 |
| None | Rizospheric soil | 3 y 4 | 19 | 41,83 |
| None | Rizospheric soil | 3 y 4 | 17 | 43,17 |
| None | Rizospheric soil | 3 y 4 | 16 | 45,13 |
| None | Rizospheric soil | 3 y 4 | 25 | 45,24 |
| None | Rizospheric soil | 3 y 4 | 21 | 49,68 |
| None | Rizospheric soil | 3 y 4 | 26 | 78,55 |
| None | Rizospheric soil | 3 y 4 | 9 | 96,00 |
